# Multi-Echo Investigations of Positive and Negative CBF and Concomitant BOLD Changes

**DOI:** 10.1101/2022.09.05.506629

**Authors:** Ratnamanjuri Devi, Jöran Lepsien, Kathrin Lorenz, Torsten Schlumm, Toralf Mildner, Harald E. Möller

**Affiliations:** Max Planck Institute for Human Cognitive and Brain Sciences, Leipzig, Germany

**Author notes:** These authors contributed equally (Toralf Mildner, Harald E. Möller).

## Abstract

Unlike the positive blood oxygenation level-dependent (BOLD) response (PBR), commonly taken as an indication of an ‘activated’ brain region, the physiological origin of negative BOLD signal changes (i.e. a negative BOLD response, NBR), also referred to as ‘deactivation’ is still being debated. In this work, an attempt was made to gain a better understanding of the underlying mechanism by obtaining a comprehensive measure of the contributing cerebral blood flow (CBF) and its relationship to the NBR in the human visual cortex, in comparison to a simultaneously induced PBR in surrounding visual regions. To overcome the low signal-to-noise ratio (SNR) of CBF measurements, a newly developed multi-echo version of a center-out echo planar-imaging (EPI) readout was employed with pseudo-continuous arterial spin labeling (pCASL). It achieved very short echo and inter-echo times and facilitated a simultaneous detection of functional CBF and BOLD changes at 3 T with improved sensitivity. Evaluations of the absolute and relative changes of CBF and the effective transverse relaxation rate, 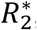, the coupling ratios, and their dependence on CBF at rest, CBF_rest_, indicated differences between activated and deactivated regions. Analysis of the shape of the respective functional responses also revealed faster negative responses with more pronounced post-stimulus transients. Resulting differences in the flow-metabolism coupling ratios were further examined for potential distinctions in the underlying neuronal contributions.

**Highlights:**

- Introduction of multi-echo center-out EPI for investigating concomitant CBF and BOLD changes in regions of positive (PBR) and negative BOLD response (NBR).
- ΔCBF timecourses closely follow those of 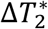 with negative signals exhibiting faster responses and more pronounced post-stimulus transients.
- Decreases in CBF appear to warrant a larger change in NBR than CBF increases in PBR regions.
- Consideration of baseline CBF values is important in comparisons of relative coupling ratios (δ*s*_BOLD_/δcbf) between brain regions.
- Discussion of potential excitatory and inhibitory neuronal feed forward control of CBF and CMRO_2_ in PBR and NBR.

## 1 Introduction

It is well established in functional magnetic resonance imaging (fMRI) that a regional cerebral blood flow (CBF) increase is the main physiological contributor to the positive blood oxygenation level-dependent (BOLD) response (PBR). More controversial is the nature of a sustained BOLD signal decrease from baseline, the negative BOLD response (NBR), which can be evoked by a suitable stimulation paradigm. Varying and often conflicting reports on its vascular and metabolic contributors have hindered the identification of its physiological basis. As a result, discussions of the NBR have considered a variety of possible mechanisms, including vascular blood steal (Harel et al., 2002; Hu and Huang, 2015; Kannurpatti and Biswal, 2004; Ma et al., 2017; Puckett et al., 2014), venous back pressure (Bandettini, 2012; Goense et al., 2012; Huber et al., 2014; Shmuel et al., 2006), neuronal activation (Devor et al., 2008; Nagaoka et al., 2006; Schridde et al., 2008; Shih et al., 2009), or inhibition (Boorman et al., 2010; Huber et al., 2014; Logothetis, 2002; Mullinger et al., 2014; Nakata et al., 2019; Shmuel et al., 2006, 2002; Wilson et al., 2019). While differing modalities [fMRI, positron emission tomography (PET), optical imaging], contrast mechanisms [iron oxide nanoparticles, vascular space occupancy, arterial spin labeling (ASL)], experimental setups, and biological conditions (humans, awake and anesthetized animals, primates or non-primates) could explain some of the discrepancies, it is also possible for the NBR to be structurally dependent (Gouws et al., 2014; Huber et al., 2019; Jorge et al., 2018; Provencher et al., 2018; Shih et al., 2009) resulting in regional and, to some extent, inter-subject variabilities.

Generally, the outcome of models describing the relationship of CBF, the cerebral blood volume (CBV), and the cerebral metabolic rate of oxygen consumption (CMRO_2_) and their contributions to the BOLD response (Buxton, 2021, 2012; Buxton et al., 2004) not only depends on a variety of parameters but also on the experimental error of the input data (Guidi et al., 2020, 2016; Huber et al., 2019). An accurate measurement of the CBF response obtained simultaneously with the BOLD signal could aid in a more quantitative understanding of the NBR, especially in the human visual cortex for which an inhibitory basis of the NBR is reasonably well established (Boillat et al., 2020; Martínez-Maestro et al., 2019; Shmuel et al., 2002; Wilson et al., 2019).

Non-invasive ASL techniques provide the best alternative to radionuclide-based or dynamic susceptibility contrast-enhanced CBF measurements. However, ASL suffers from an inherently low signal-to-noise ratio (SNR) and contamination by the BOLD signal at commonly used echo times (TEs). This is further complicated in regions of an NBR due to weaker effect sizes and higher variability compared to that of the PBR. Nevertheless, CBF measurements have not only been used to determine their contribution (Huber et al., 2014; Wilson et al., 2019) but also to estimate measures of CMRO_2_ using calibrated BOLD approaches (Davis et al., 1998; Hoge et al., 1999) in NBR regions (Fukunaga et al., 2008; Mullinger et al., 2014; Pasley et al., 2007; Stefanovic et al., 2005, 2004).

In our study performed at 3 T, we employed pseudo-continuous ASL (pCASL) (Dai et al., 2008) to achieve a high SNR albeit at the cost of a somewhat lower temporal resolution. A previously recommended set of parameters (Alsop et al., 2015; Lorenz et al., 2018) was used, and labeling duration and post-labeling delay (PLD) were optimized by estimating the arterial transit time in the visual cortex in combination with an ASL model (Alsop and Detre, 1996; Mildner et al., 2014; Wang et al., 2002). A further means to improve the SNR was the combination with a segmented echo planar-imaging (EPI) readout (Chapman et al., 1987), Double shot EPI with Center-out Trajectories and Intrinsic NaviGation (DEPICTING) (Hetzer et al., 2011), supporting acquisitions at TE<2 ms, where BOLD contamination was negligible (Devi et al., 2019). As an extension of this preliminary work, we introduced a multi-echo (ME) version of the sequence, ME-DEPICTING, which allows for the simultaneous measurement of CBF and BOLD responses. In one substudy, ME-DEPICTING and ME-EPI were, therefore, compared in PBR regions to examine their general sensitivities for detecting CBF and BOLD responses (“*substudy 1*”). Another substudy focused on a quantification of changes of CBF and the effective transverse relaxation rate, 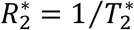, in regions of PBR and NBR as well as an investigation of derived quantities, coupling ratios, or dependencies on baseline values (“*substudy 2*”). Additionally, to derive hypotheses on the underlying neuronal mechanisms based on the available experimental data, CMRO_2_ values were estimated and speculations about feed-forward neuronal control of the CBF and CMRO_2_ were discussed through implementation and extension of the Wilson-Cowan model (Wilson & Cowan, 1972) proposed recently by (Buxton, 2021).

## 2 Materials & Methods

### 2.1 Participants

Eighteen healthy volunteers (30±7 years, 8 female) gave written informed consent before undergoing the experiments that had been approved by the Ethics Committee at the Medical Faculty of Leipzig University. The first 13 subjects (30±8 years, 6 female) took part in *substudy 1* as well as *substudy 2*, while the remaining five participated only in *substudy 2*. All subjects were right-handed and had normal to corrected vision.

### 2.2 Functional Paradigm

A small 8-Hz flickering radial checkerboard (Supplementary Figure S1), subtending a visual angle of approximately 1.8° was used for visual stimulation (Huber et al., 2014; Wade and Rowland, 2010). It is known to induce a sustained NBR in the primary visual cortex (V1). Each functional cycle lasted 20 repetitions, starting with a rest block, which consisted of a blank gray screen of 12 repetitions (42 s), followed by the task block of 8 repetitions (28 s). The paradigm was programmed using Presentation (v17.2, Neurobehavioral Systems, Berkeley, CA, USA). A central, colored fixation point was present throughout the experiment. Subjects were instructed to focus on this dot and press a button whenever it changed color. Their attention was monitored by visually tracking their responses. A post-scan questionnaire was provided for a self-evaluation of their performance and comfort-level rating of the stimulus.

### 2.3 Magnetic Resonance Acquisitions

Experiments for *substudy 1* were conducted on a 3T MAGNETOM Prisma^fit^ scanner (Siemens Healthineers, Erlangen, Germany) using a 32-channel receive head coil. Functional CBF changes were measured by labeling arterial blood in a plane 65 mm caudal from the nasal root for a duration of 1,500 ms using an optimized pCASL radiofrequency (RF) pulse train with an average gradient amplitude, *G*_av_=0.65 mT/m; average RF pulse amplitude, *B*_1,av_=1.5 μT; RF pulse length of 600 μs; and 50% RF duty cycle (Lorenz et al., 2018). Following a PLD of 1200 ms, 12 slices (nominal resolution 3×3×4 mm^3^, 0.8mm slice gap, field of view 192 mm, matrix 64×64, bandwidth 2230 Hz/Px) were acquired along the calcarine sulcus. The two readout modules had the following specifications:

- ME-EPI: TE_1_ / TE_2_ / TE3 = 8ms/21.2ms/34.4ms; repetition time, TR =3,500 ms; GRAPPA (GeneRalized Autocalibrating Partial Parallel Acquisition) factor 2; partial-Fourier factor 6/8.
- ME-DEPICTING: TE_1_ / TE_2_ / TE_3_ =1.7ms/10.7ms/19.7ms; TR =3,552 ms; GRAPPA factor 2.

The schematic of the ME-DEPICTING sequence and the corresponding k-space trajectory are given in Figure 1. The first k-space tiles for all echoes are sampled before their second tiles. This is realized by gradient rewinders after the respective phase blips and achieves both an ultra-short TE and a reduced inter-echo time (ΔTE). Supplementary Figure S2 gives the expanded sequence diagram for all three echoes. For an inter-segment intensity correction (Hetzer et al., 2011), a C++ functor was implemented in the Siemens Image Calculation Environment (ICE). To avoid double contours in DEPICTING acquisitions, a correction of off-resonance effects is typically required by applying the information of local magnetic field offset values, Δ*B*_0_ (Hetzer et al., 2011; Patzig et al., 2021). In the current study, this correction could be omitted due to the short image acquisition time, *T*_acq_=16 ms, and the small offsets: Δ*B*_0_ maps showed that the shift (i.e., *T*_acq_×Δ*B*_0_) was less than half the nominal voxel size in about 99% of all voxels in the visual cortex. The quality of the raw images of both sequences can be assessed through Supplementary Figure S3.

**Figure 1.**
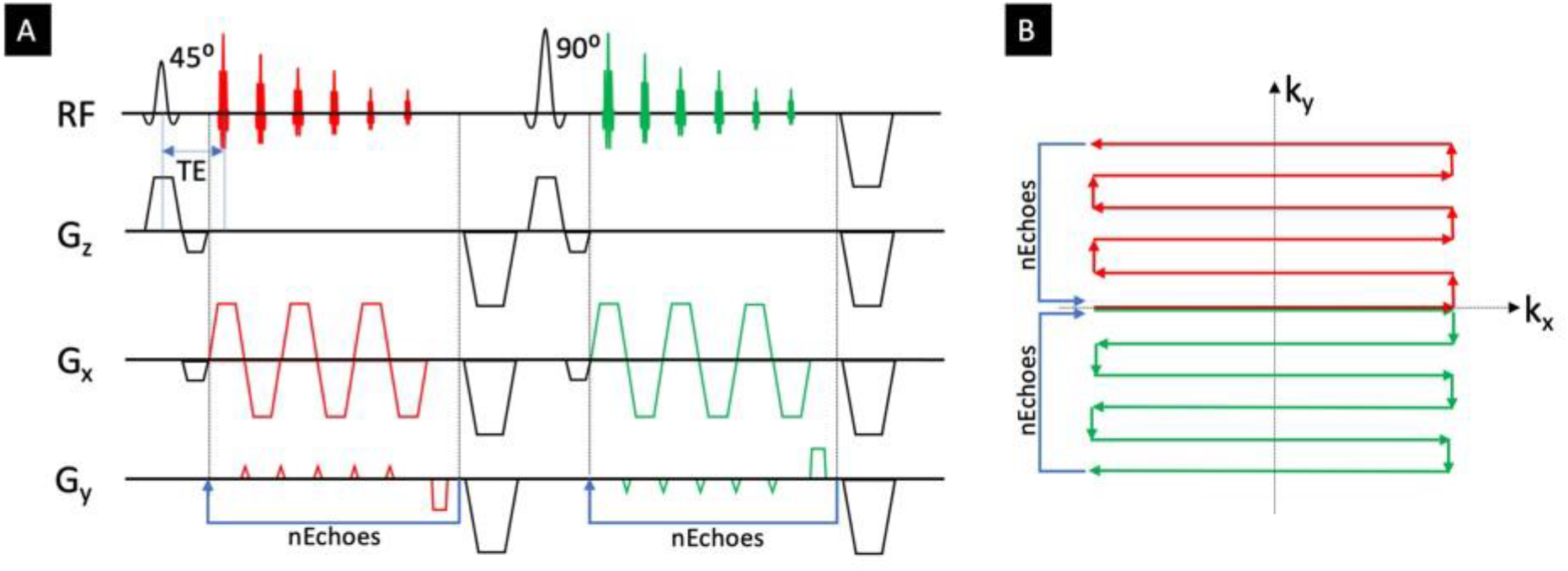
Schematic of the ME-DEPICTING sequence **(A)** along with the corresponding k-space trajectory **(B)**. The loop extension for each segment is chosen according to the number of echoes, n. Note that another flyback gradient lobe without data acquisition has to be added along the read direction in case of an odd number of measured lines per segment.

Experiments for *substudy 2*, on the other hand, were conducted on two 3T scanners: the MAGNETOM Prisma^fit^ and a MAGNETOM Skyra^fit^ (Siemens Healthineers, Erlangen, Germany). A 32-channel receive head coil was used in both cases. The pCASL parameters were identical to *substudy 1*, with the exception that the labeling plane was taken either at the base of the cerebellum or at a plane 65mm caudal to the nasal root. Similarly, 10-12 slices (nominal resolution 3×3×4 mm^3^, 0–0.8mm slice gap, field of view 192 mm, matrix 64×64, bandwidth 2230 Hz/Px) were acquired along the calcarine sulcus using TR=3,500–3,552 ms and otherwise identical ME-DEPICTING specifications as in *substudy 1*.

Functional runs of *substudy 1* comprised of five functional cycles (50 control/label pairs) while those of *substudy 2* using pCASL-prepared ME-DEPICTING consisted of ten functional cycles (100 control/label pairs) and lasted approximately 12 min. In the 13 subjects who participated in both studies, data were acquired within the same session. A break of 15–20 min without stimulation was placed between the acquisitions of the two substudies. Additionally, the order of the ME-EPI and ME-DEPICTING acquisitions was shuffled to avoid primacy and recency effects. Auxiliary resting-state runs of 5–6min duration were also recorded with the pCASL-prepared readouts in three subjects. Two volumes without pCASL preparation were acquired before resting-state or functional acquisitions.

Three-dimensional (3D) *T*_1_ -weighted anatomical references were available from separate sessions with Magnetization-Prepared RApid Gradient Echo (MP-RAGE)(Mugler and Brookeman, 1990) or Magnetization-Prepared 2 Rapid Acquisition Gradient Echoes (MP2RAGE) (Marques et al., 2010) acquisitions with previously published parameters (Streitbürger et al., 2014). For registration purposes, two 2D gradient-recalled echo (GRE) scans (1.5mm nominal in-plane resolution; TE =3 ms; TR =150 ms; flip angle 60°) were obtained during each session, one at the start and the other at the end, with identical slice geometry as the functional scans.

Subjects scanned with the Prisma^fit^ scanner are preceded by a ‘P’ before numbers denoting the order in which they were acquired, while those scanned with the Skyra^fit^ start with an ‘S’. Subject number 7 was scanned twice, once with each scanner and is, hence, referred to as ‘P7’ and ‘S7’ within the manuscript.

### 2.4 Data Preprocessing and Analysis

Data were preprocessed and analyzed using Statistical Parametric Mapping (SPM12; Wellcome Trust Centre for Neuroimaging, UCL, London, UK) implemented in MATLAB 2019b (Mathworks, Natick, MA, USA) and additional scripts written in Interactive Data Language (IDL 8.1, Exelis Visual Information Solutions, Boulder, CO, USA).

All functional data were preprocessed in an identical manner. The time series of the first echo was realigned to its first repetition, and the resulting transformation parameters of each repetition were used to resample the time series of all echoes. For spatial normalization to the Montreal Neurological Institute (MNI) coordinate space (Evans et al., 1993), a co-registration procedure capable of improving the realignment quality for small numbers of slices was performed. The individual MP2RAGE/MP-RAGE volume was first registered on the structural 2D GRE images, which were then co-registered with the first functional EPI data. The resulting transformation was applied to create a warped version of the structural 3D image. The warped 3D image was then used to normalize the functional time series of all echoes to MNI space with an output resolution of 2 mm. For this procedure, the GRE scan acquired with the shorter delay to the respective functional run was selected. The normalized time series were finally temporally high-pass filtered at a cut-off frequency equal to two functional cycles [1/(40 TR)] and 3D-Gaussian filtered at 2mm full width at half maximum.

Using the IDL function LINFIT, “derived” time series of the model parameters 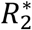 and the signal intensity at zero echo time, *S*_0_, were extracted from the preprocessed ME image volume *n* acquired at echo time TE_*n*_ with voxel intensities *S*_*n*_ via linear regression of the expression In 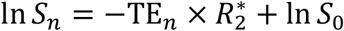. A hybrid time series of signal intensities *S*_sum_ obtained by combining the multiple echoes by a weighted summation was also generated (Poser et al., 2006; Posse et al., 1999). The weights were computed from the echo times and the expected BOLD contrast according to the fitted 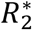 of each voxel (see Eq. 6 of Ref. 52).

For statistical analysis, an ASL-specific general linear model (GLM) (Hernandez-Garcia et al., 2010; Mumford et al., 2006) was implemented in IDL. Contrary to the original model, the regressor for the baseline ASL signal was shifted to a range between 0 and −1 instead of 0.5 and −0.5. This choice tangibly removes the dependence of the concomitantly derived BOLD contrast from the ASL contrast by making the BOLD regressor correspond only to the fluctuations of ASL control images rather than to that of the mean of the ASL control/label images. Furthermore, estimated realignment parameters were included as nuisance regressors. The GLM was then applied to the time series of TE_1_ and *S*_sum_, wherein the ASL and BOLD regressors were respectively taken to be the covariates-of-interest. The model was also applied to the fitted time series of *S*_0_ and 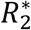 to obtain additional quantitative ASL and BOLD contrasts, respectively. These were used for comparison with the main contrasts obtained using TE_1_ and *S*_sum_ in the sequence comparison (*substudy 1*), while only that of 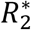 was used to quantify BOLD signal changes in *substudy 2*.

The baseline ASL contrast of the GLM (difference between baseline and label images in the resting period) and the sum of both baseline and functional ASL contrasts (difference between baseline and label images in the stimulation period) were scaled by the baseline contrast (control signal in the resting period). Then, they were converted to CBF at rest (CBF_rest_) and ΔCBF = CBF_stim_ − CBF_rest_ (CBF_stim_ is CBF during stimulation) in units of ml/100g/min, respectively, by using a two-compartment model (Alsop and Detre, 1996; Mildner et al., 2014; Wang et al., 2002).The following model parameters were assumed (Lorenz et al., 2018): brain-blood partition coefficient 0.9 ml/g, gray-matter (GM) density 1.04 g/ml, pCASL inversion efficiency 90%, arterial blood and tissue relaxation times *T*_1_ and 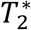 of 1,664 ms and 1,330 ms, and of 50 ms and 55 ms, respectively. The model also included assumed values of the arterial and the tissue transit time of 1,000 ms and 1,600 ms, respectively. The BOLD signal was quantified either in terms of the relative signal change, δ*S*_BOLD_ = ((*S*_stim_ − *S*_rest_)/*S*_rest_) × 100 (obtained from *S*_sum_), or in terms of the quantitative rate change 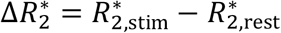. The latter value was taken directly from the corresponding *β*_*i*_ parameter of the GLM fit to the 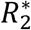 time series.

#### 2.4.1 Comparison of ME-DEPICTING and ME-EPI (*Substudy 1*)

Results from the first-level analysis were thresholded at a Bonferroni-corrected significance level of *p* <0.05 to obtain CBF and BOLD activation maps. A visual-cortex mask was additionally applied to these maps. This mask was generated by combining V1 and extrastriate areas V2, V3, V4 and V5 from the Jülich Histological Atlas (Amunts et al., 2000; Malikovic et al., 2007; Rottschy et al., 2007) in FSLeyes of the FMRIB Software Library (FSL) (Jenkinson et al., 2012) and transforming the result to the MNI space. Potential differences due to intra-session motion were compensated by multiplying this visual-cortex mask by a brain mask common to the corresponding scans for the individual subject. The FSL Brain Extraction Tool (BET) (Smith, 2002) was used to generate the individual brain masks. Supra-threshold voxels and statistical significance levels obtained with ME-EPI and ME-DEPICTING were then evaluated within this region. A measure of inter-subject variation was obtained by counting the number of occurrences of a particular voxel across the significantly activated regions of all subjects, thus yielding maps of consistently activated voxels for the ASL and BOLD contrasts and each sequence. Additionally, the numbers of activated voxels appearing in at least one subject and those activated consistently in 6 or more subjects were determined.

The sensitivities of both sequences for BOLD- and CBF-based activation measurements were also compared in terms of the more reliable contrast-to-noise ratio (CNR), which is known to be less dependent of statistical thresholds and the number of time points (Geissler et al., 2007). The significant regions obtained from the first-level analysis were used to generate two separate masks for each subject: one based on common regions of CBF activation (“CBF-mask”) obtained from the *S*_0_ and TE_1_ contrasts of both sequences, and the other one based on common regions of BOLD activation (“BOLD-mask”) obtained with *S*_sum_ and 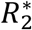. The CNR was then assessed twice: *(i)* in the smaller region defined by the CBF-mask, and *(ii)* in the larger region defined by the BOLD-mask. The CNR over the respective region was obtained by dividing the mean of the beta image estimated in the GLM fit by the standard deviation of the residual noise.

Additionally, the temporal stability was evaluated from the auxiliary resting-state scans by considering the temporal SNR (tSNR) of the pairwise subtracted ASL time series (TE_1_ and *S*_0_ data) and pairwise averaged BOLD time series (*S*_sum_ and 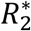 data) within the whole slab as well as in the visual-cortex mask described above. Both masks were multiplied by a GM mask of >50% probability, obtained from SPM12 segmentations of the respective MNI-warped anatomical scans.

#### 2.4.2 Relationship of ΔCBF and 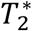 (*Substudy 2*)

Regions of significant BOLD signal change were identified by thresholding the corresponding statistical maps of *S*_sum_ at *p*<0.05 (Bonferroni-corrected). Similarly, regions of significant CBF changes were obtained by thresholding the statistical maps of the TE_1_ data at *p*<0.01. False positives arising from potential spurious contributions from large pial veins and cerebrospinal fluid (CSF) were additionally controlled in both maps by masking out voxels with 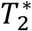 values at rest, 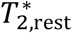, outside a range between 25 and 60 ms (Wansapura et al., 1999). Similarly, voxels exhibiting CBF_rest_ values outside an acceptable range between 20 and 120 ml/100g/min were excluded. Further analysis for each subject was then restricted to overlapping regions of the surviving significant CBF and BOLD activation [“positive” region-of-interest (ROI)] and deactivation (“negative” ROI). The following quantities were evaluated within the ROIs: δ*S*_BOLD_ (in %), ΔCBF (in ml/100g/min), the relative CBF change (δcbf = ΔCBF/CBF_rest_ in %), 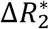 (in s^−1^), CBF (in ml/100g/min), and 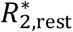 and 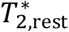 (in s^−1^ and ms, respectively). Mean absolute errors over the ROI were calculated from the mean relative standard errors of the corresponding GLM contrast, StdErr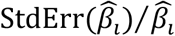, multiplied by the corresponding mean value over the ROI in absolute or percent units. Ratios of concomitant changes of the functional quantities or ‘coupling ratios’, expressed as 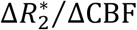 and δ*S*_BOLD_ /δcbf, were also determined along with their maximum errors, propagated from the mean absolute errors over the ROI of the contributing parameters.

The following timecourses were extracted from the ROIs: *(i)* CBF timecourses generated by pairwise subtraction of the TE_1_ data [(*S*_control_ − *S*_label_)/*S*_control_]*×* 100, followed by quantification using the two-compartment model. *(ii)* 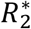 timecourses of corresponding temporal resolution by taking the running average of labeling and control time points of the 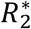 data 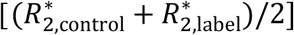; *(iii)* 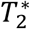 timecourses computed from *(ii)* for a more intuitive comparison with the CBF results. Cycle-averaged and interpolated ΔCBF and 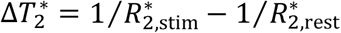 timecourses were further used to evaluate temporal characteristics, such as the time to peak of the primary signal (TTP_1_) and that of the post-stimulus transients (TTP_2_).

Paired two-tailed Student’s *t*-tests were used to assess differences in the mean values of all quantities between the two ROIs.

#### 2.4.3 CMRO_2_ estimation (*Substudy 2*)

Relative changes in CMRO_2_ (δcmro_2_ in %) within the two ROIs were also estimated using the Davis model (Davis et al., 1998; Hoge et al., 1999), such that,

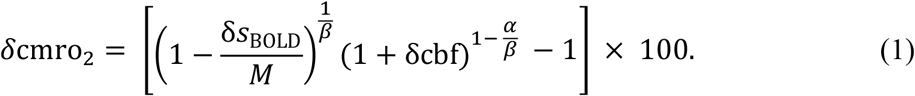

Owing to the absence of gas calibration experiments in the present study, the BOLD calibration constant, *M* was assumed to be 4% and was estimated for our TE_eff_ of ∼14 ms from the reported value of ∼8% at TE = 29 ms (Hare and Bulte, 2016; Whittaker et al., 2016). The model parameters for both ROIs were then assumed to be the same. The Grubbs coefficient, *α* was assumed to be 0.2 (Chen and Pike, 2009), while two values of the exponent *β* of the Davis model were used: *β*=1.5 (Davis et al., 1998; Hoge et al., 1999; Uludag et al., 2004), and the more commonly used *β*=1.3 at 3T (Chiarelli et al., 2007; Mark et al., 2011). The maximum errors of the estimated δcmro_2_ were propagated from the mean absolute errors over the ROI of the measured parameters. The flow-metabolism coupling ratio, *n* = δcbf/δcmro_2_ was then evaluated for the two ROIs.

Furthermore, the potential feed-forward excitatory and inhibitory control of CBF and CMRO_2_ as investigated in (Buxton, 2021) through the implementation of the Wilson-Cowan model (Wilson and Cowan, 1972), was replicated and tested for its fit to our data. This model is referred to here as the Wilson-Cowan flow-metabolism model. It follows the assumption of relative changes in CMRO_2_ driven mainly by excitatory activity (δcmro_2|WC_ ∼ *E*) and those of CBF brought about by a combination of excitatory and inhibitory activity (δcbf_WC_ ∼ *E* + 𝒳_PBR_ × *I*; 𝒳 describes the relative weighting of the inhibitory activity, which was assumed to be 𝒳_PBR_ = 1.5 for the PBR). The relation between δcbf and δcmro_2_ in the positive ROI was examined in the manner described in Supplementary C of (Buxton, 2021). To summarize, we used a simplified version of the Wilson Cowan model:

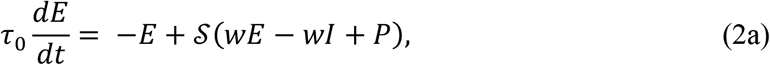

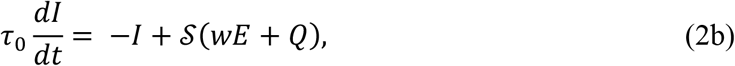

where *t* is time, τ_0_ is a decay time constant (here set to 10 ms), 𝒮(*x*) = [1 + *e*^−*a*(𝒳−θ)^]^−1^ − (1 + *e*^*a*θ^)^−1^ is a sigmoidal transfer function of gain *a* and threshold θ, *w* the strength of the synaptic connections, and *P* and *Q* the external excitatory and inhibitory inputs, respectively. The *E* and *I* values corresponding to *P* ranging from 0 to 2.5 at a constant *Q* of 0.2 were scaled and equated to relative changes in CBF and CMRO_2_. The proportionality constant for both quantities was obtained by equating the excitatory activity, *E* corresponding to maximum external excitatory input, *P* to 30% δcmro_2_ that is, 30/*E*(*P* = 2.5) ≈ 39. The negative signs, −*E* and −*I* represent the leakage in the absence of an input current; this input is represented by the parameters within the sigmoid function, such that the excitatory current is *wE* − *wI* + *P* and the inhibitory current is *wE* + *Q*.

Assuming a similar neuronal control in regions of NBR, the model was adapted here to the negative ROI. By building on the assumption of a resting initial condition of almost zero activity, that is, *E*(0) = 0 and *I*(0) = 0, a decrease in the input current would support a deactivation. Negative values *E* and *I* were found possible only if a decreasing sigmoid function is considered, which corresponds to the following equations:

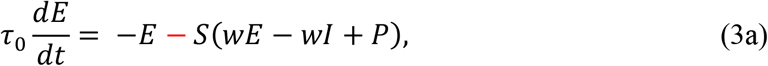

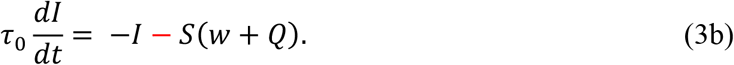

We have to note that a negative current (or firing rate) is not physiologically possible and the above equations work only with the assumption of a baseline condition (Sadeghi et al., 2020). Another issue that needs further evaluation is that of the sign of leakage. Two sets of these equations were, hence, implemented, one with a positive leak and the other with a negative leak. The same values of the remaining parameters were assumed as for activation, with the exception of *Q*. The model was tested for a constant *Q* = 0.2 as well as *P* = *Q* linearly increasing from 0 to 2.5. The resulting *E*(*P* = 2.5) was then equated to a δcmro_2_ of –30% and the proportionality constant applied to obtain δcbf_WC_ ∼ *E* + 𝒳_NBR_ × *I*; 𝒳_NBR_ = 1.5.

Additionally, the δcmro_2_ values of the model (δcmro_2|WC_) for both PBR and NBR were converted to δ*S*_BOLD|WC_ by applying the Davis model with the same assumptions as in the CMRO_2_ estimation stated above, to generate the Wilson-Cowan δ*S*_BOLD_/δcbf model. This was done to test the adherence of this model with our experimentally obtained δ*S*_BOLD_ and δcbf values (see also Supplementary Methods).

## 3 Results

### 3.1 Comparison of ME-DEPICTING and ME-EPI (*Substudy 1*)

#### 3.1.1 Sensitivity to Functional Changes

Robust and significant CBF and BOLD activations were obtained in all scans of 12 subjects. Subject P6 had to be omitted from this substudy due to a rotational-motion artefact (pitch >0.8°) in the EPI scan. Figure 2 presents functional maps, and Figure 3 shows examples of cycle-averaged pCASL timecourses for both sequences, which were extracted from a common region of significant CBF activation. The PBR is visible in the timecourses through an increase of the control signal intensity (orange solid lines) during the stimulation (shaded area). Notably, the very weak BOLD response obtained with ME-DEPICTING at TE_1_ =1.7 ms (top left) is reflected by an almost flat control signal. The stronger BOLD contamination accompanied by increased signal fluctuations in the ME-EPI timecourses is also visible.

**Figure 2.**
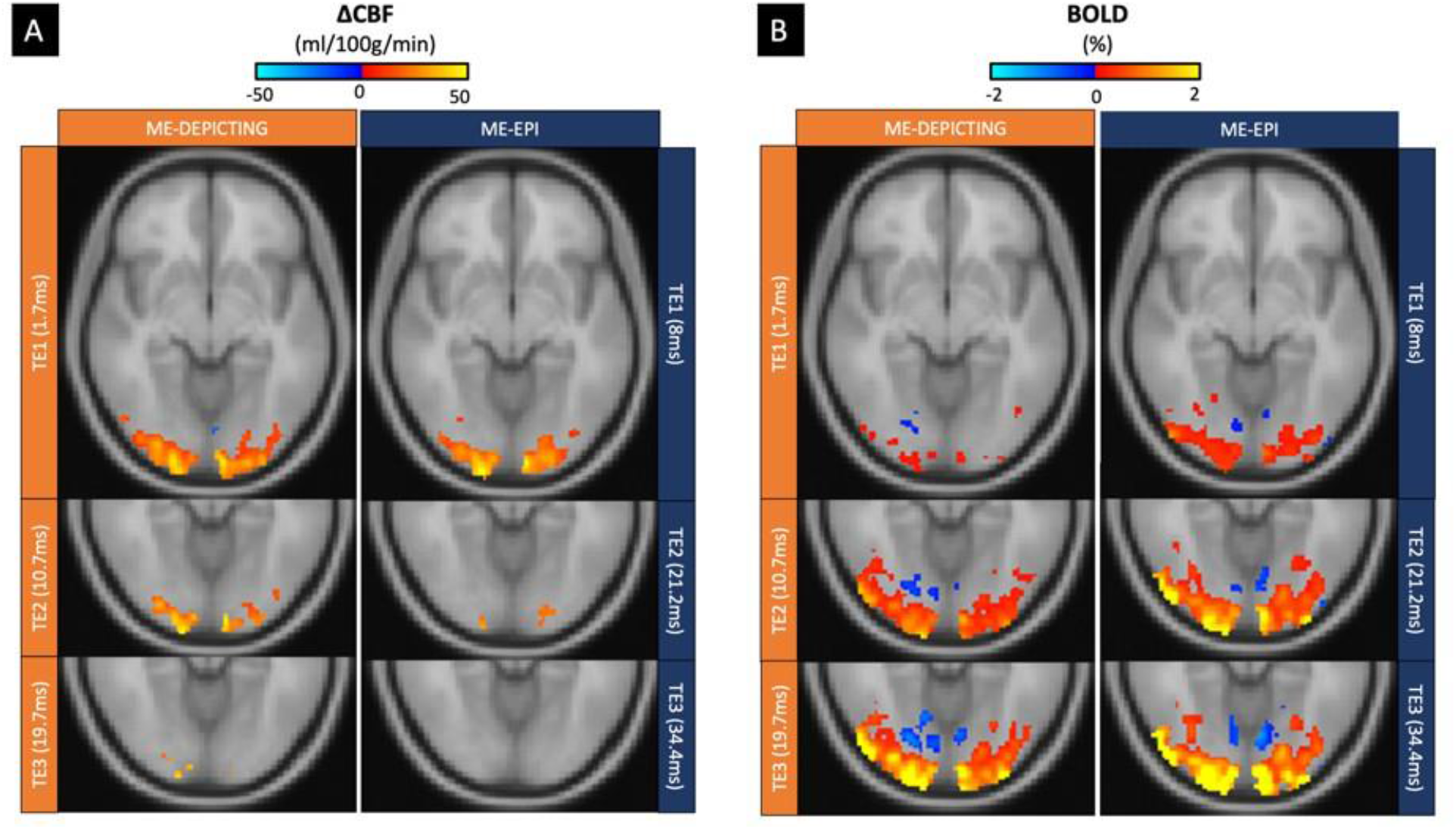
Subject-level ΔCBF **(A)** and BOLD functional maps **(B)** obtained with all three echoes of ME-DEPICTING and ME-EPI acquisitions in the same subject. At TE_3_ (bottom row), regions of both PBR (red-to-yellow color gradient) and NBR (blue color gradient) are detected with both sequences, decreasing in size at shorter TE due to the reduced BOLD response (B). In the PBR regions, an activation-induced increase in CBF is evident at TE_1_ (top row), which diminishes at longer TE because the progressively increased BOLD signal (of opposite sign) interferes with the ASL signal. Note that the functional runs of substudy 1 as shown here were always acquired after substudy 2 with only half the number of cycles (i.e. reduced sensitivity) and generally weakened responses, probably indicating expectation or repetition suppression (Summerfield et al., 2008). Therefore, only the stronger and, hence, more robust PBR was selected for the quantitative sequence comparison.

**Figure 3.**
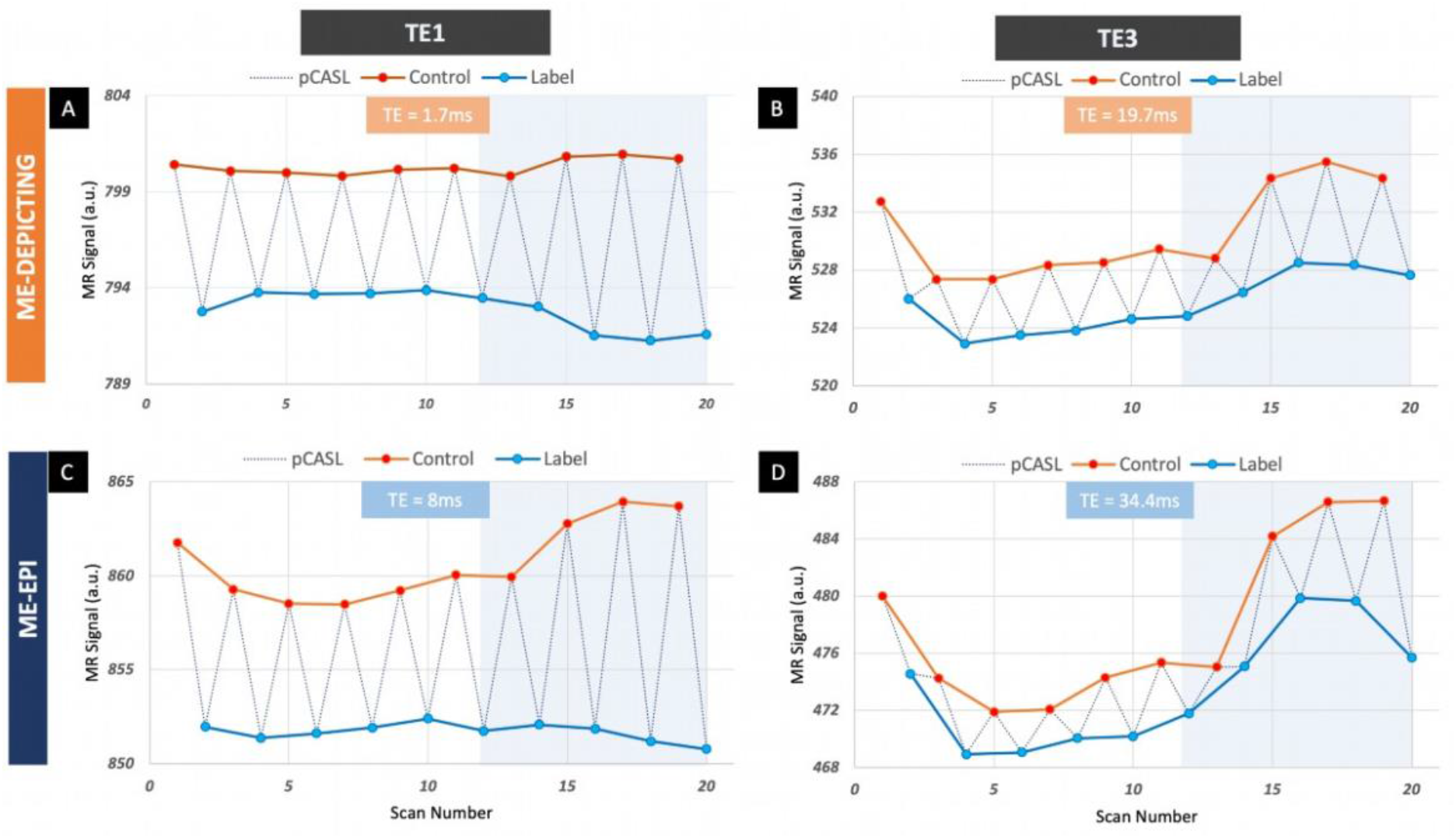
Cycle-averaged pre-processed pCASL timecourses in a representative subject, obtained at the shortest (TE_1_) and longest echo time (TE_3_) with ME-DEPICTING **(A, B)** and ME-EPI **(C, D)**. All timecourses were extracted from a common region of significant stimulus-induced CBF change based on the images acquired at TE_1_. The shaded region indicates the duration of the visual task.

The number of supra-threshold voxels for CBF and BOLD activation and the average *p*-values, ⟨*p*⟩, obtained from the TE_1_ and *S*_sum_ data are compared in Figure 4. For ME-DEPICTING, the average extent of CBF activation significantly (*p*=0.001) exceeded that of ME-EPI by 43% (ME-DEPICTING: 1,539±870 voxels; ME-EPI: 1,075±809 voxels). A superior statistical significance, and hence, sensitivity for measuring CBF changes with ME-DEPICTING is also highlighted by the consistently smaller *p*-values in all subjects. The average extent of BOLD activation was similar for both sequences (ME-DEPICTING: 4,309±1,293 voxels; ME-EPI: 4,400±1,629 voxels). Interestingly, the average *p* -value of BOLD activation for ME-DEPICTING was still smaller by approximately 20%.

**Figure 4.**
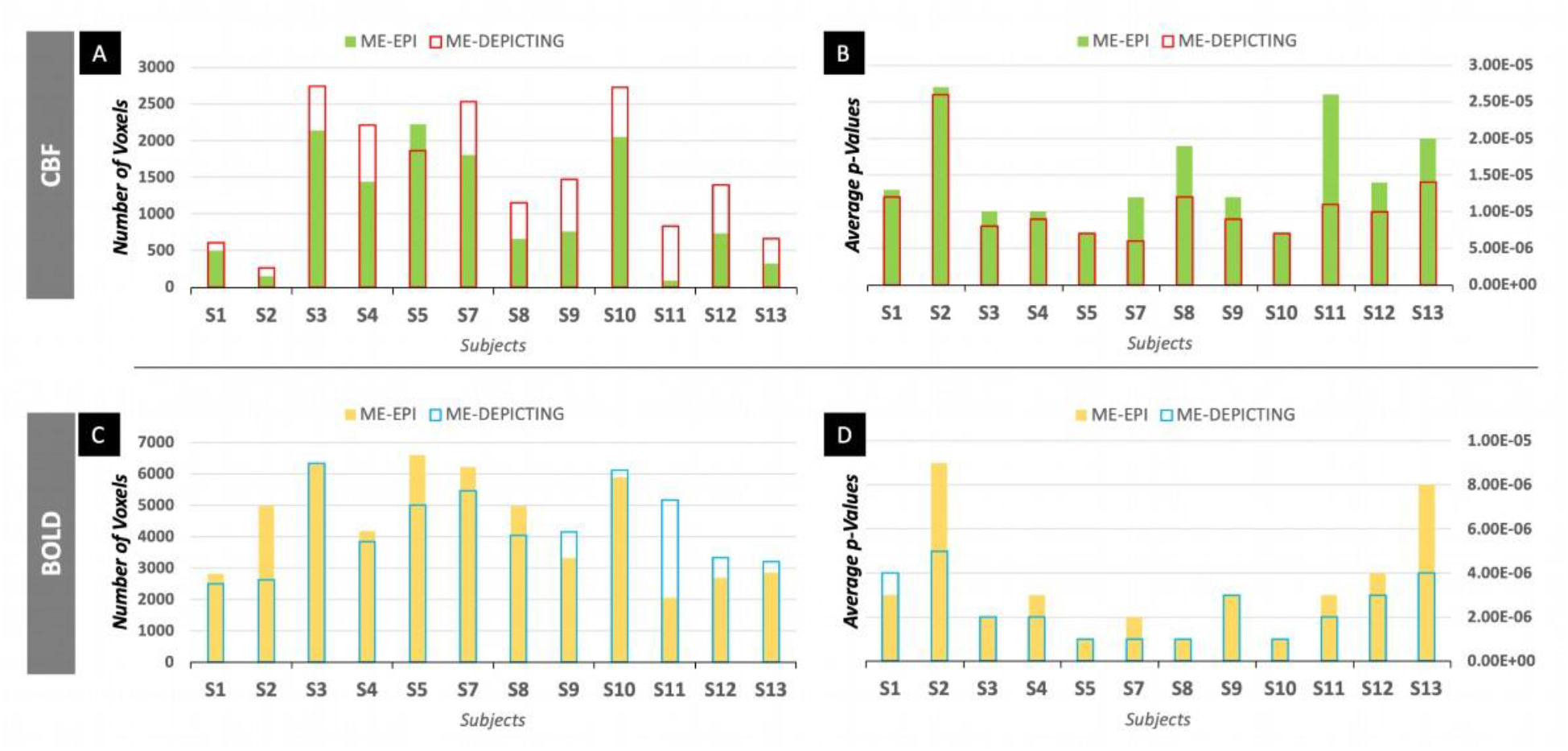
Differences in sensitivity for measuring stimulus-induced CBF **(A, B)** and BOLD **(C, D)** signal changes with ME-DEPICTING and ME-EPI in 12 subjects. Both the number of supra-threshold voxels **(A, C)** and average p-values **(B, D)** are shown. The values were obtained from the TE_1_ contrast for the CBF **(A, B)** and from the S_sum_ contrast for the BOLD measurements **(C, D)**.

This trend was also observed for the consistently activated voxels (Figure 5): In the CBF measurement, 5,717 voxels with counts ≥1 and a mean count of 3.24 were obtained with ME-DEPICTING compared to 5,252 voxels and a mean count of 2.45 with ME-EPI. Furthermore, 19.4% and 9.5% of these voxels had ≥6 counts for ME-DEPICTING and ME-EPI, respectively. Although fewer voxels with counts ≥1 were obtained in the BOLD-activation map with ME-DEPICTING (11,548 vs. 13,271), the number of voxels with counts ≥6 was still larger (3,926 vs. 3,635). Figure 6A gives a graphical representation of these results.

**Figure 5.**
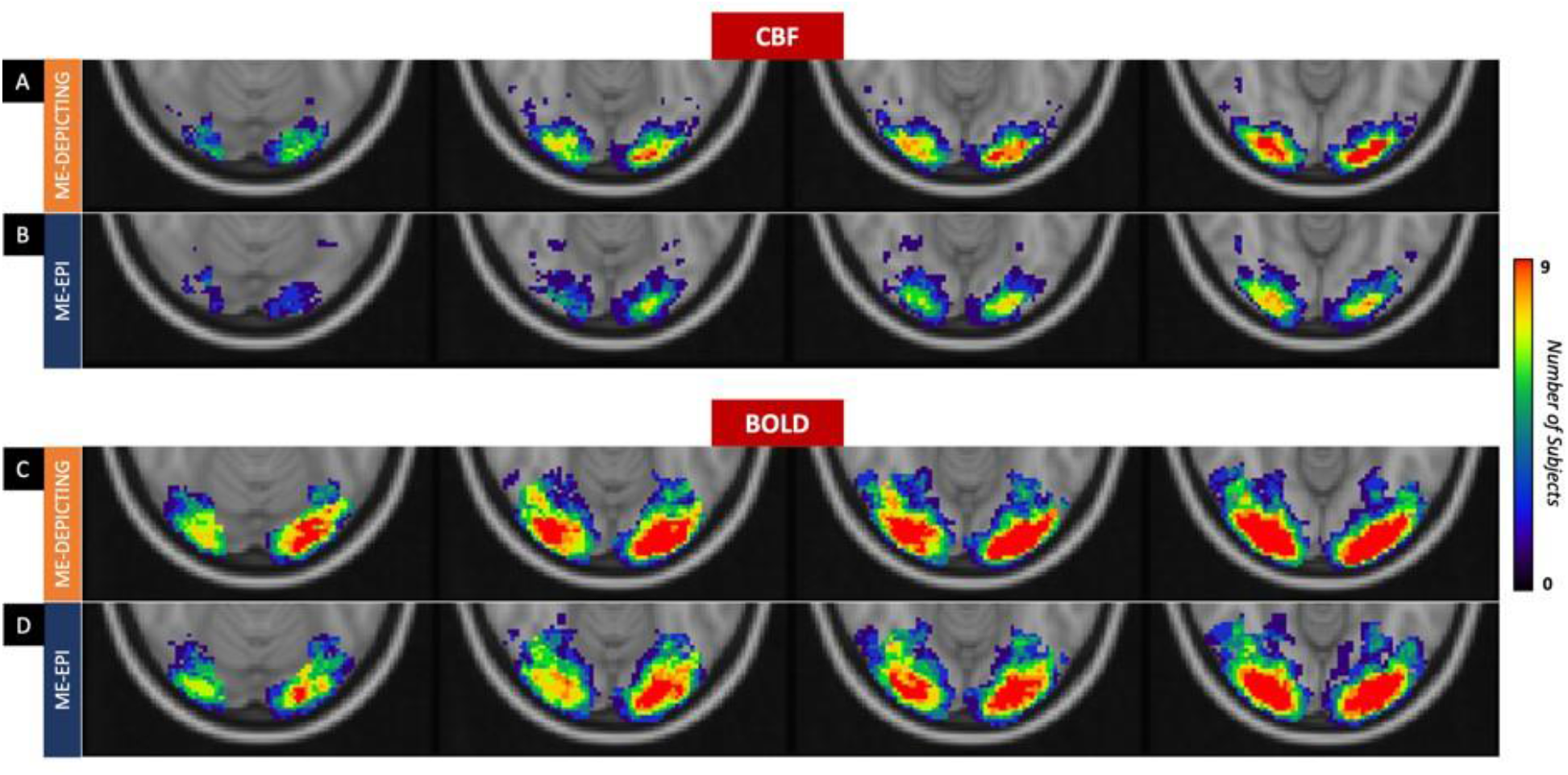
Maps showing voxels activated across all subjects for stimulus-induced positive CBF **(A, B)** and BOLD signal changes **(C, D)** obtained with ME-DEPICTING **(A, C)** and ME-EPI **(B, D)**. Each voxel bears a color that represents the number of occurrences of the particular voxel across our study sample. Note that regions of NBR were not considered in this analysis.

**Figure 6.**
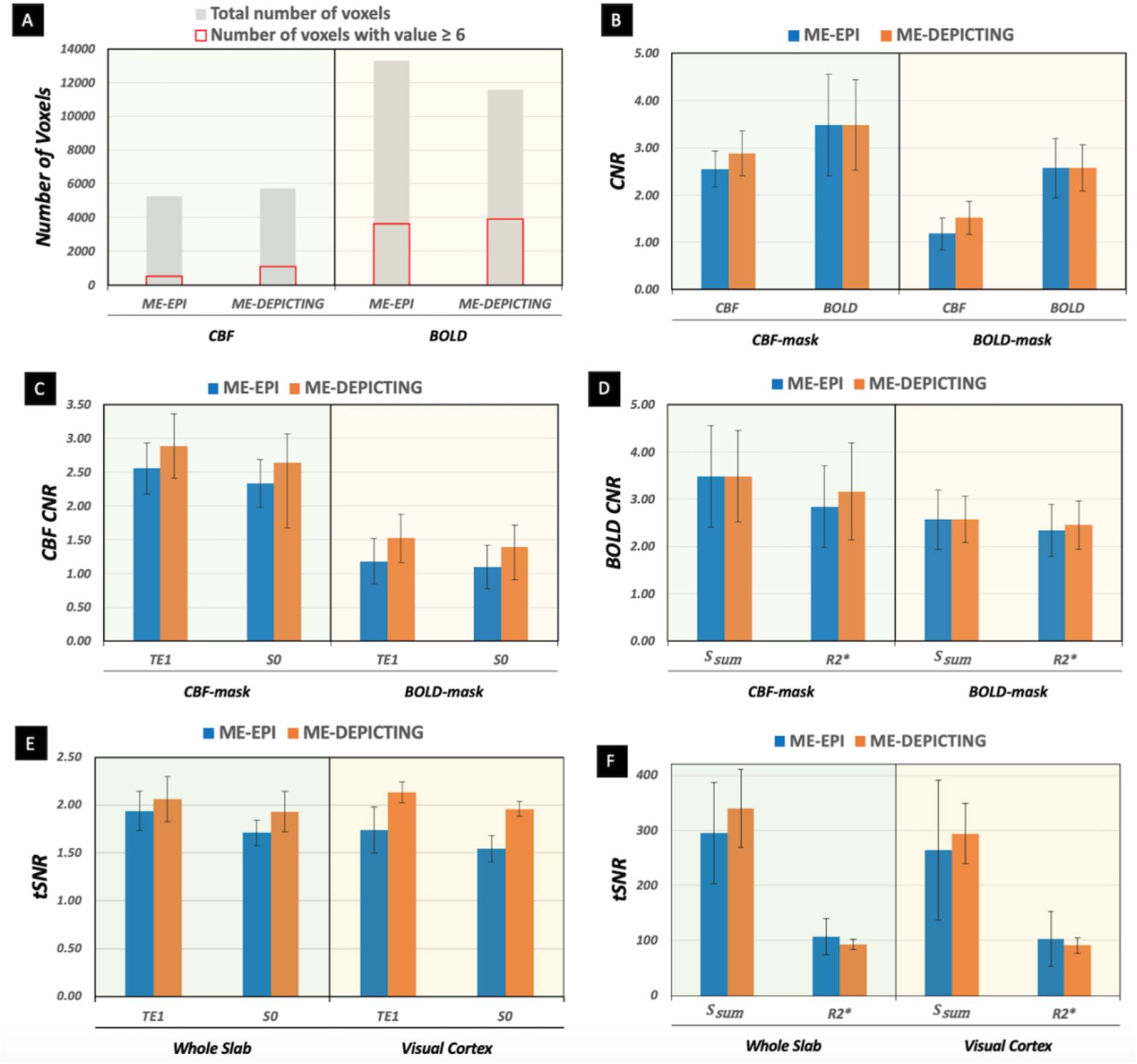
Comparison of ME-EPI and ME-DEPICTING group-averaged results. **(A)** Total number of voxels (solid gray bars) with counts ≥1 of consistent activation in 12 subjects along with the number of most commonly activated voxels (counts ≥6; open red bars). **(B)** CNR values (means and standard deviations from 12 subjects) for stimulus-induced positive CBF and BOLD signal changes obtained within the CBF-mask **(left)** and within the BOLD-mask **(right)**. CBF-based **(C)** and BOLD-based **(D)** CNR obtained with TE_1_ and S_sum_ as seen in **(B)** compared with derived contrasts S_0_ and 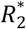, respectively. Results obtained with ME-EPI and ME-DEPICTING are shown as solid blue and orange bars, respectively. tSNR values for ASL **(E)** and BOLD **(F)** data in three subjects averaged over the whole slab **(left)** and averaged over the visual cortex **(right)**.

The results of these first- and second-level analyses were also confirmed by the CNR evaluation with subject-averaged values of 2.89±0.48 and 2.55±0.38 for ME-DEPICTING and ME-EPI in the CBF-mask, as well as 1.52±0.35 and 1.18±0.34 in the BOLD-mask, respectively (Figure 6B). Interestingly, the CNR improvement for CBF detection with ME-DEPICTING was higher for the larger BOLD-mask (29%, *p*=0.001) than for the CBF-mask (13%, *p*=0.01). By contrast, the CNRs for BOLD-effect detection with ME-DEPICTING and ME-EPI were similar (3.48±1.07 and 3.48±0.96 in the CBF-mask and 2.57±0.63 and 2.58±0.49 in the BOLD-mask, respectively).

#### 3.1.2 Derived Contrasts and tSNR of Resting ASL Data

The statistical power of presumably purer CBF and BOLD contrasts obtained from the *S*_0_ and 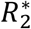 data, respectively, was compared to routinely obtained results from TE and *S* data. For ΔCBF from *S*_0_ compared to TE_1_, the average size of the activated region significantly decreased (*p*<0.05) by 19% and 12% for ME-EPI and ME-DEPICTING, respectively, with a concomitant increase of ⟨*p*⟩ by 20% and 7%. For the BOLD response obtained from 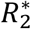 compared to *S*_sum_, the average size of the region of significant BOLD-based activation decreased by 13% and 3% for ME-EPI and ME-DEPICTING, respectively. A greater stability of routinely used contrasts was also suggested by counts of consistently activated voxels: *S*_0_ compared to TE_1_ data yielded 11% and 6% fewer voxels with counts ≥1 for ME-EPI and ME-DEPICTING, respectively, and 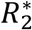 compared to *S*_sum_ data had 8% and 1.3% fewer voxels.

For both sequences, the CNR of CBF increases was reduced by 8−9% for *S*_0_ compared to TE_1_ in both masks (Figure 6C). Interestingly, the drop in BOLD-CNR (Figure 6D) for 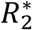 instead of *S*_sum_ roughly doubled for ME-EPI yielding 23% and 10% in the CBF- and BOLD-mask, respectively, compared to 10% and 5% for ME-DEPICTING. Figure 6E shows the comparison of tSNR taken from pairwise subtracted ASL resting-state data yielding 1.74±0.24 and 2.13±0.11 with TE_1_in the visual-cortex mask for ME-EPI and ME-DEPICTING, respectively, as well as reductions by 11% and 8% upon using *S*_0_. A larger reduction in the tSNR was seen for both readouts with their 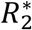 data compared to the corresponding *S*_sum_ data (Figure 6F).

To summarize, the sensitivity of time series “derived” by exponential fitting of ME data (yielding *S*_0_ and 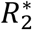) did not reach that of “direct” series (employing TE_1_ and *S*_sum_). This sensitivity loss was relatively small for ME-DEPICTING, and CBF measurements in visual cortex based on *S*_0_ from ME-DEPICTING scans still outperformed ME-EPI scans at TE_1_.

### 3.2 Relation of ΔCBF and 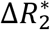 (*Substudy 2*)

#### 3.2.1 Positive vs. Negative ROI Statistics

A significant PBR and NBR was consistently identified in expected regions of the visual cortex. Corresponding regions with significant CBF increase and decrease were observed in all subjects, with the exception of S16. Its region of significant CBF decrease did not match its regions of significant NBR (Supplementary Table S1). The average spatial overlap of BOLD- and CBF-based functional maps was 90±8% for positive and 51±27% for negative responses. The positive ROIs were larger and less spatially variable than their negative counterparts. An example is shown in Figure 7.

**Figure 7.**
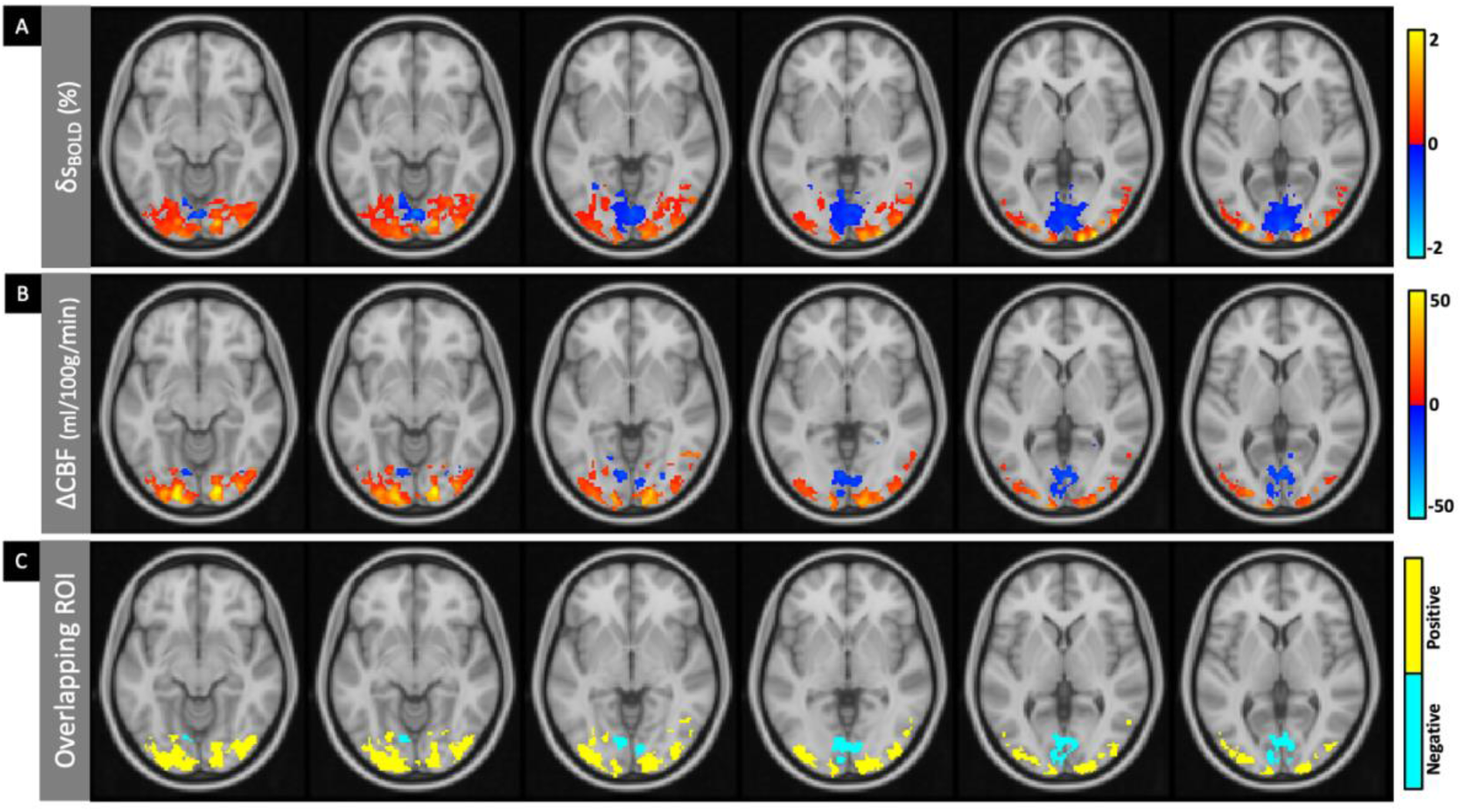
Representative spatial activity maps obtained in subject P1 from simultaneous acquisitions of percent BOLD-signal change (A) and absolute CBF change (B) overlaid on the T_1_-weighted MNI template. The overlaps of the regions of activation (90%) and deactivation (84%) obtained with both measures define corresponding positive and negative ROIs for the individual subject (C). The BOLD map was thresholded at p<0.0001 and the CBF map at p<0.01. Masks based on resting CBF and T2* values have been applied to all maps.

The size of the negative ROIs ranged from a minimum of 6 (subject P2) to a maximum of 1,712 voxels (subject S18), while the size of the positive ROIs varied between 357 (subject P14) and 4611 voxels (subject P7). Group-averaged relative changes of (0.76±0.14)% and (44.2±7.5)% were obtained in BOLD and CBF measurements, respectively, in the positive ROIs corresponding to absolute differences 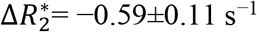 and ΔCBF=20.9±3.7 ml/100g/min. For the negative ROIs, relative changes were (−0.42±0.11)% and (−21.2±4.5)%, respectively, corresponding to 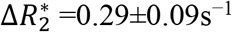 and ΔCBF = −12.9±2.1 ml/100g/min. Single-subject values of all functional contrasts including their resting-state values are summarized in Table 1. Mean positive-to-negative amplitude ratios of 2.2±0.5 and 1.6±0.3 were obtained for δcbf and ΔCBF, respectively, which were of similar size as the BOLD signal (1.8±0.3) and 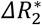 ratios (2.1±0.4).

**Table 1.** Single-subject and group-averaged values (plus/minus one standard deviation) of relative changes δS_BOLD_ and δcbf, absolute changes 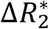 and ΔCBF, as well as resting-state values of 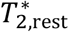 and CBF_rest_ as obtained from the positive and negative ROIs, respectively. Asterisks after subject numbers indicate female participants.

| Subjects | ROI size<br>(voxels) | | $\delta s_{\text{BOLD}}$<br>(%) | | $\Delta R_2^*$<br>($s^{-1}$) | | $T_{2,\text{rest}}^*$<br>(ms) | | $\delta \text{cbf}$<br>(%) | | $\Delta \text{CBF}$<br>(ml/100g/min) | | $\text{CBF}_{\text{rest}}$<br>(ml/100g/min) | |
| --- | --- | --- | --- | --- | --- | --- | --- | --- | --- | --- | --- | --- | --- | --- |
|  | Positive | Negative | Positive | Negative | Positive | Negative | Positive | Negative | Positive | Negative | Positive | Negative | Positive | Negative |
| P1 | 2087 | 801 | 0.61 | -0.38 | -0.48 | 0.28 | 39.2 | 42.8 | 53.3 | -18.0 | 20.3 | -11.3 | 41.6 | 68.6 |
| P2 | 1661 | 6 | 0.56 | -0.31 | -0.45 | 0.17 | 38.0 | 53.1 | 38.9 | -19.5 | 13.9 | -11.5 | 39.2 | 63.3 |
| P3* | 3728 | 1000 | 0.71 | -0.35 | -0.54 | 0.24 | 42.3 | 47.4 | 40.4 | -22.5 | 22.2 | -13.3 | 57.9 | 61.8 |
| P4* | 3109 | 235 | 0.79 | -0.37 | -0.62 | 0.23 | 40.1 | 42.3 | 47.6 | -23.3 | 24.1 | -11.7 | 56.6 | 53.8 |
| P5 | 3093 | 14 | 0.91 | -0.35 | -0.70 | 0.24 | 40.7 | 36.0 | 55.6 | -15.9 | 23.0 | -11.8 | 44.7 | 75.7 |
| P6 | 908 | 21 | 0.58 | -0.25 | -0.42 | 0.18 | 39.0 | 38.2 | 46.5 | -26.8 | 17.7 | -11.6 | 40.8 | 43.6 |
| P7* | 4611 | 703 | 0.71 | -0.36 | -0.57 | 0.28 | 39.3 | 41.3 | 31.9 | -14.4 | 21.3 | -10.6 | 66.8 | 74.7 |
| P8 | 2486 | 232 | 0.84 | -0.52 | -0.66 | 0.36 | 39.1 | 41.9 | 49.3 | -23.2 | 17.5 | -12.3 | 37.8 | 54.8 |
| P9* | 2932 | 272 | 0.74 | -0.52 | -0.57 | 0.31 | 40.5 | 44.9 | 40.1 | -16.1 | 23.0 | -12.3 | 59.3 | 77.2 |
| P10* | 2628 | 543 | 0.59 | -0.31 | -0.45 | 0.24 | 42.9 | 48.7 | 31.2 | -15.1 | 20.7 | -12.5 | 69.1 | 85.1 |
| P11 | 1681 | 47 | 0.95 | -0.54 | -0.70 | 0.36 | 37.5 | 38.5 | 53.7 | -28.6 | 21.9 | -15.4 | 44.5 | 56.4 |
| P12 | 2732 | 72 | 0.78 | -0.43 | -0.62 | 0.32 | 40.3 | 32.3 | 52.3 | -20.9 | 25.8 | -17.6 | 52.7 | 84.2 |
| P13* | 1993 | 365 | 0.73 | -0.45 | -0.59 | 0.32 | 42.5 | 46.6 | 43.4 | -20.0 | 23.1 | -14.7 | 55.5 | 74.7 |
| P14* | 357 | 101 | 0.62 | -0.34 | -0.51 | 0.25 | 40.8 | 44.6 | 38.3 | -17.9 | 13.3 | -10.0 | 38.2 | 56.3 |
| 15 | 1266 | 64 | 0.73 | -0.46 | -0.51 | 0.29 | 43.1 | 46.7 | 41.1 | -29.1 | 17.1 | -13.8 | 44.8 | 47.9 |
| S16 | 2226 | 0 | 0.94 | - | -0.76 | - | 38.5 | - | 53.5 | - | 22.7 | - | 45.8 | - |
| *S17\** | 2715 | 170 | 0.67 | -0.40 | -0.53 | 0.22 | 37.8 | 45.8 | 36.4 | -22.8 | 19.9 | -12.2 | 59.0 | 56.1 |
| *S7\** | 3590 | 427 | 0.99 | -0.56 | -0.80 | 0.43 | 35.9 | 39.1 | 43.7 | -21.9 | 22.4 | -13.2 | 53.6 | 62.5 |
| *S18* | 1843 | 1712 | 0.94 | -0.70 | -0.71 | 0.54 | 42.3 | 43.9 | 54.0 | -25.8 | 27.8 | -16.8 | 55.5 | 67.5 |
| **Mean ±** | **2402 ±** | **357 ±** | **0.76 ±** | **-0.42 ±** | **-0.59 ±** | **0.29 ±** | **40.0 ±** | **43.0 ±** | **44.2 ±** | **-21.2 ±** | **20.9 ±** | **-12.9 ±** | **50.7 ±** | **64.7 ±** |
| **SD** | **1026** | **440** | **0.14** | **0.11** | **0.11** | **0.09** | **2.0** | **5.0** | **7.5** | **4.5** | **3.7** | **2.1** | **9.6** | **12.1** |

Absolute signal changes plotted against CBF_rest_ are shown in Figure 8. Correlations with CBF_rest_ for the negative ROI yielded very small and insignificant coefficients of determination, *r*_2_, consistent with the assumption of an independence of 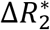 or ΔCBF from CBF_rest_. A similar behavior was observed for 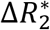 data over the positive ROI. Positive ΔCBF values, on the other hand, with a pronounced linear increase for CBF_rest_ values below ∼50 ml/min/100g, demonstrated a significant linear correlation (*p* = 0.02, *r*^2^ =0.30) with CBF_rest_ (Figure 8B). The corresponding plots of δ*S*_BOLD_ and δcbf over CBF_rest_ are provided in Supplementary Figure S4. The CBF_rest_ was additionally found to be significantly higher in the negative ROI (located entirely in V1) than in the positive ROI (located in striate and extrastriate visual areas) across all subjects (64.7±12.1 vs. 50.7±9.6 ml/100g/min; *p*<0.001). Differences in 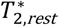 over the two ROIs were also found to be significant (43.0±5.0 vs. 40.0±2.0 ms; *p*=0.02).

**Figure 8.**
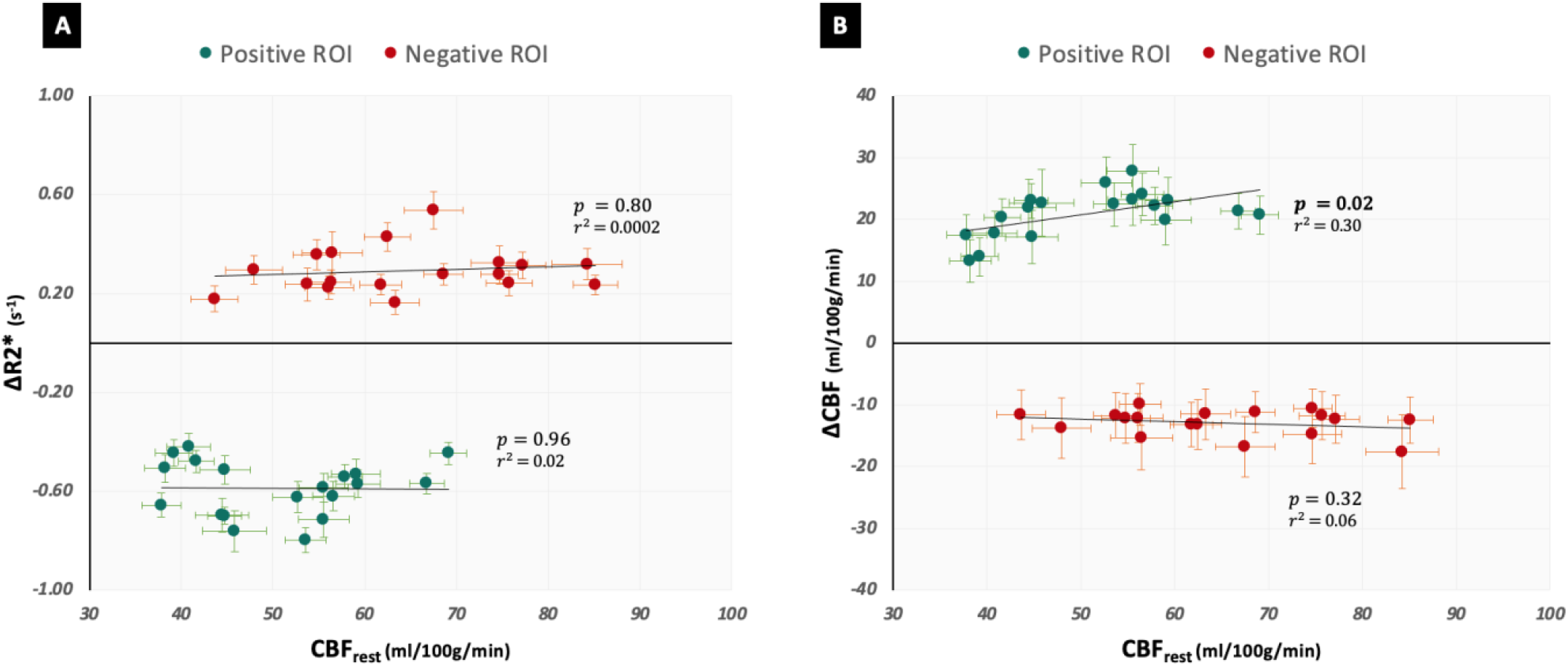
Absolute stimulus-induced changes of 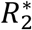 in s^−1^ **(A)** and CBF in ml/100g/min **(B)** plotted as a function of their corresponding CBF_rest_ for all subjects. Green and red circles show average values in the positive and negative ROIs, respectively. Error bars indicate the standard errors (across the ROI) of 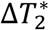, ΔCBF and CBF_rest_.

Figure 9A plots the mean δ*S*_BOLD_ against the corresponding δcbf of the two ROIs for all subjects. A very significant (*p*=10^−27^, *r*_2_=0.96) linear slope is indicated over the full range of δcbf values. Although intuitive, this linear model was found to be similar to one with a non-zero intercept [analysis of variance (ANOVA) model comparison; *F*(1,35)=3.92, *p*=0.06]. Linear fits of the separate ROIs, as denoted by the dotted lines, also revealed significant δ*S*_BOLD_/δcbf δ*S*_BOLD_/δcbf slopes which were slightly higher over the negative ROI (*p*=0.23). The individual coupling ratios over the negative ROI averaged to 0.020±0.005 and were significantly higher (*p*=0.006) than the corresponding ratios over the positive ROI which averaged at 0.017±0.003. The 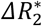 ratios were also found to be different between the two ROIs [–0.029±0.005 s^−1^/(ml/100g/min) over the positive ROI and –0.023±0.005 s^− 1^/(ml/100g/min). The relative and absolute coupling ratios for all subjects can be found in Supplementary Table S2.

**Figure 9.**
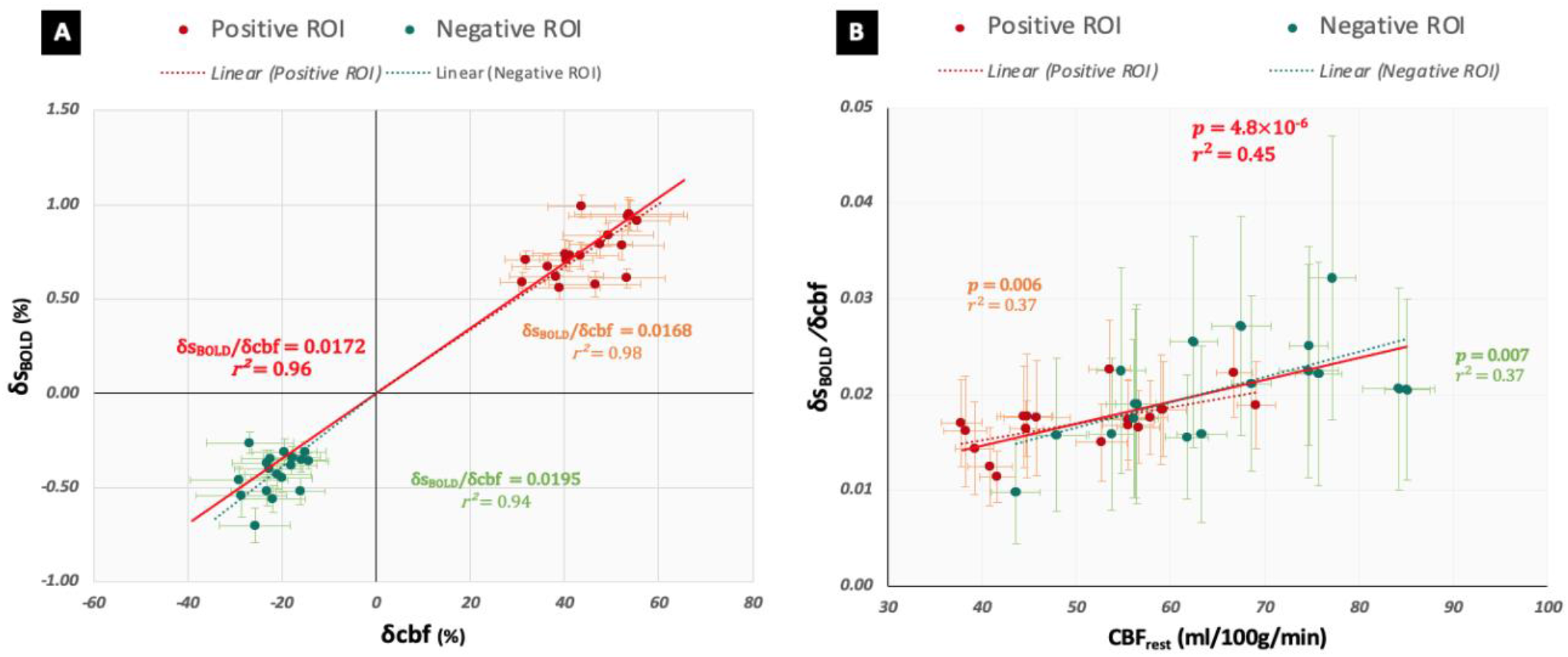
Plots of δS_BOLD_ versus δcbf over both ROIs **(A)** and δS_BOLD_/δcbf plotted as a function of CBF_rest_ **(B)** in all 18 subjects. Error bars indicate the standard errors (across the ROI) of δS_BOLD_, δcbf and CBF_rest_, respectively, while vertical error bars of **(B**) represent maximum errors propagated from standard errors of the contributing parameters of the ratios. The red solid line indicates the linear fit over all the values.

Remarkably, the δ*S*_BOLD_/δcbf coupling ratios were also found to exhibit a strong overall dependence on CBF_rest_ (*p* =4.8×10^−6^, *r*_2_ =0.45) (Figure 9B). This dependence was also significant over the individual positive (*p*=0.006, *r*_2_=0.37) as well as the negative (*p*=0.007, *r*_2_=0.37) ROIs.

#### 3.2.2 Timecourses

Individual and group-averaged ΔCBF and 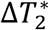 timecourses for twelve subjects are shown in Figure 10. In order to reduce noise in the derived shape parameters, only subjects with a minimum of 100 voxels in their negative ROIs were included in this analysis (*n*=12). All timecourses exhibited primary positive (PBR) or negative (NBR) peaks in response to the stimulus before returning to the baseline after its cessation. These initial returns were followed by opposite post-stimulus transient signals of weaker amplitude than the primary peaks. Moreover, the post-stimulus transients of ΔCBF were consistently less pronounced than those of their 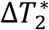 counterparts. Similar characteristics can also be observed in the timecourses of the relative δ*S*_BOLD_ and δcbf signals (Supplementary Figure S5).

**Figure 10.**
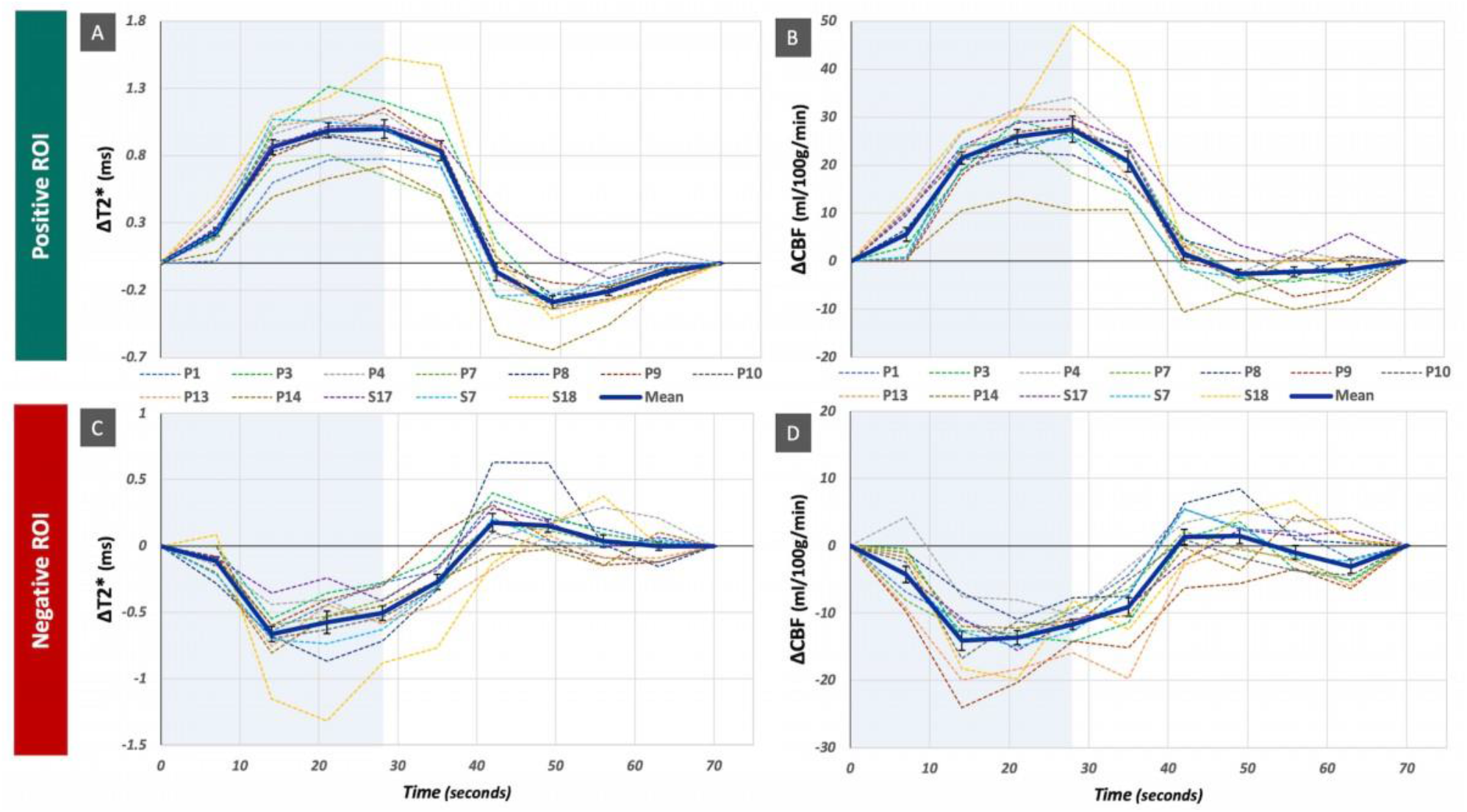
Cycle-averaged 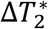 **(A, C)** and ΔCBF **(B, D)** timecourses in twelve subjects with a minimum ROI size of 100 voxels in positive **(A, B)** and negative ROIs **(C, D)** ROIs, namely P1, P3, P4, P7, P8, P9, P10, P13, P14, S17, S7 and S18. Subject-averaged mean timecourses are shown as bold solid blue lines. The error bars indicate one standard error of the mean (SEM). Blue shaded regions indicate the duration of the task.

Figure 11A depicts subject-averaged normalized 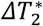 and ΔCBF timecourses for the positive and negative ROIs indicating a high level of similarity of both types of transients. This general similarity of the shapes further manifests in similar TTPs of the primary signals: Average TTP_1_ values calculated from the interpolated signals were 25.2±6.0 s and 24.3±5.4 s for the 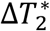 and the ΔCBF transients in the positive and 18.4±5.0 s and 20.2±5.5 s in the negative ROI, respectively. These results also indicate that the TTP_1_ intervals of the negative ROI are shorter than in the positive ROI. This is further highlighted in Figure 11B, which compares normalized transients of positive and inverted transients of negative ROIs for both 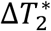 and ΔCBF.

**Figure 11.**
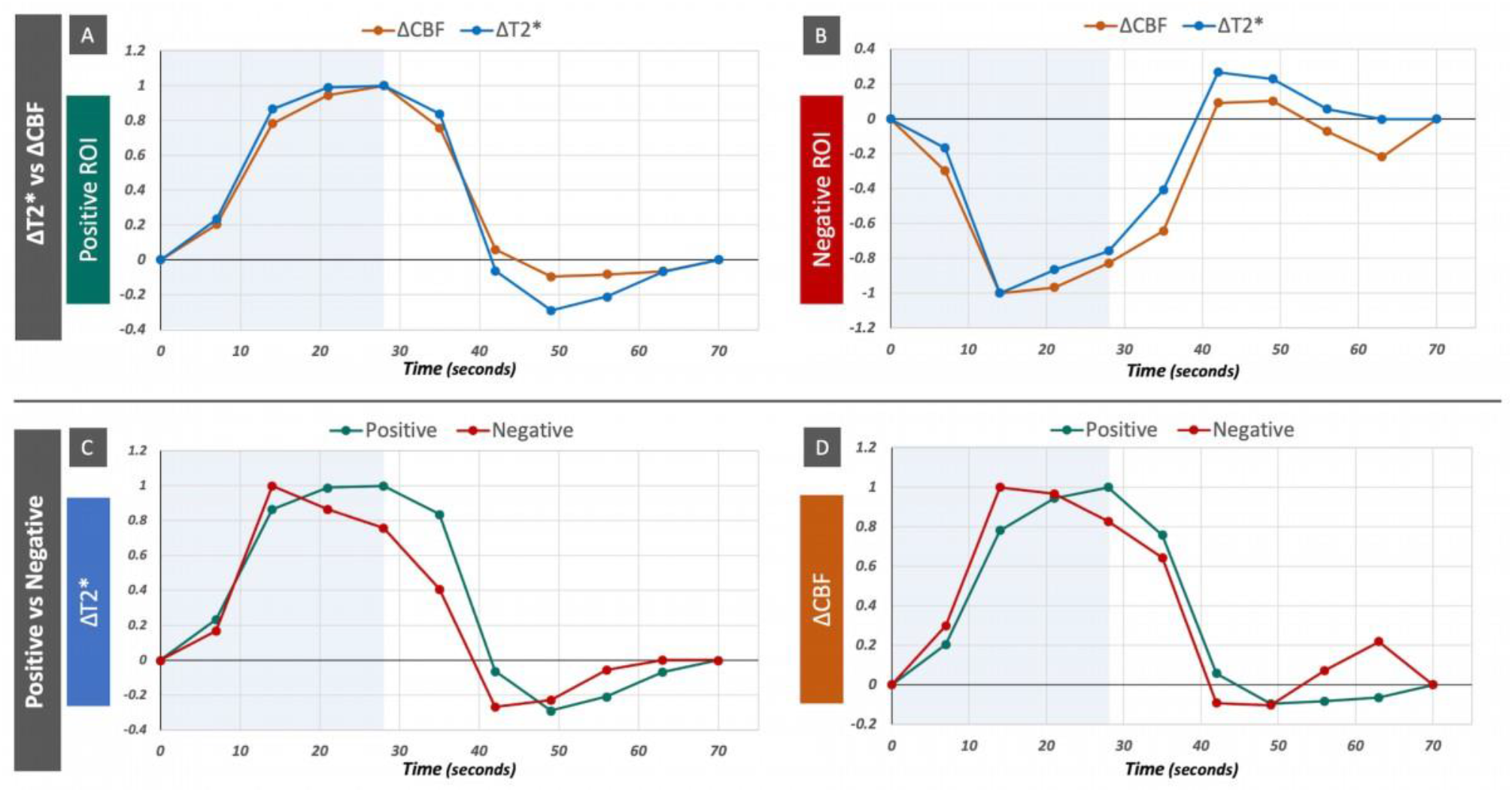
Normalized mean positive and negative transients as indicated by bold solid blue lines in Figure 10 from the same 12 subjects. Comparison between mean 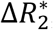 and ΔCBF timecourses in the positive **(A)** and negative ROIs **(B)** as well as comparison between mean positive and negative 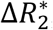 **(C)** and ΔCBF timecourses **(D)**. Blue shaded regions indicate the duration of the task.

Further similarities between 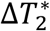 and ΔCBF were observed for delays TTP_2_ of the post-stimulus undershoot (PSU) in the positive and overshoots (PSO) in the negative ROI with average PSU values of 47.9±3.2 s and 49.3±4.5 s for 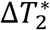 and ΔCBF, respectively, and average PSO values of 47.7±7.4 s and 44.6±4.8 s. We note that all TTP_2_ values refer to the first peak after stimulus cessation as a transient post-stimulus oscillatory behavior was observed in some cases. Furthermore, the PSOs in the negative ROI appeared to be stronger than the PSUs in the positive ROI with amplitude ratios of the post-stimulus and the primary peak of 0.48 and 0.34 for 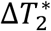 in the negative and positive ROI, respectively, and corresponding values of 0.31 and 0.19 for ΔCBF.

### 3.3 Flow-metabolism coupling (*Substudy 2*)

δcmro_2_ averaged to (19.7±4.5)% and (–13.1±4.0)% over the positive and negative ROI, respectively, for *β*=1.5. A similar positive:negative ratio of ∼1.5 was obtained for δcmro_2_ estimated with *β*=1.3 as well, averaging at (16.2±4.3)% and (–11.7±4.0)%. Figure 12 plots the mean δcbf over the corresponding δcmro_2_ estimated with the two *β* values, for all subjects in a manner similar to Figure 9A. Complementary to the δ*S*_BOLD_/δcbf coupling ratios, the subject-averaged flow-metabolism coupling ratio, *n*, based on the measured δcbf and the estimated δcmro_2_ was found to be significantly different between the two ROIs (2.3 vs.1.7 for δcmro_2_ estimated with *β*=1.5; *p*=2.3×10^−7^ and 2.9 vs. 1.9 for *β*=1.3; *p*=2.5×10^−6^). The strength of the linear correlation was slightly higher for the negative ROIs, and the overall the coupling was found to be tighter for values estimated with *β*=1.5 (*r*_2_= 0.96 vs *r*_2_= 0.94). Contrary to the slopes of δ*S*BOLD/δcbf, the individual fitted slopes, δcbf/δcmro_2_ remained significantly (*p*=5.4×10^−4^) different between the two ROIs.

**Figure 12.**
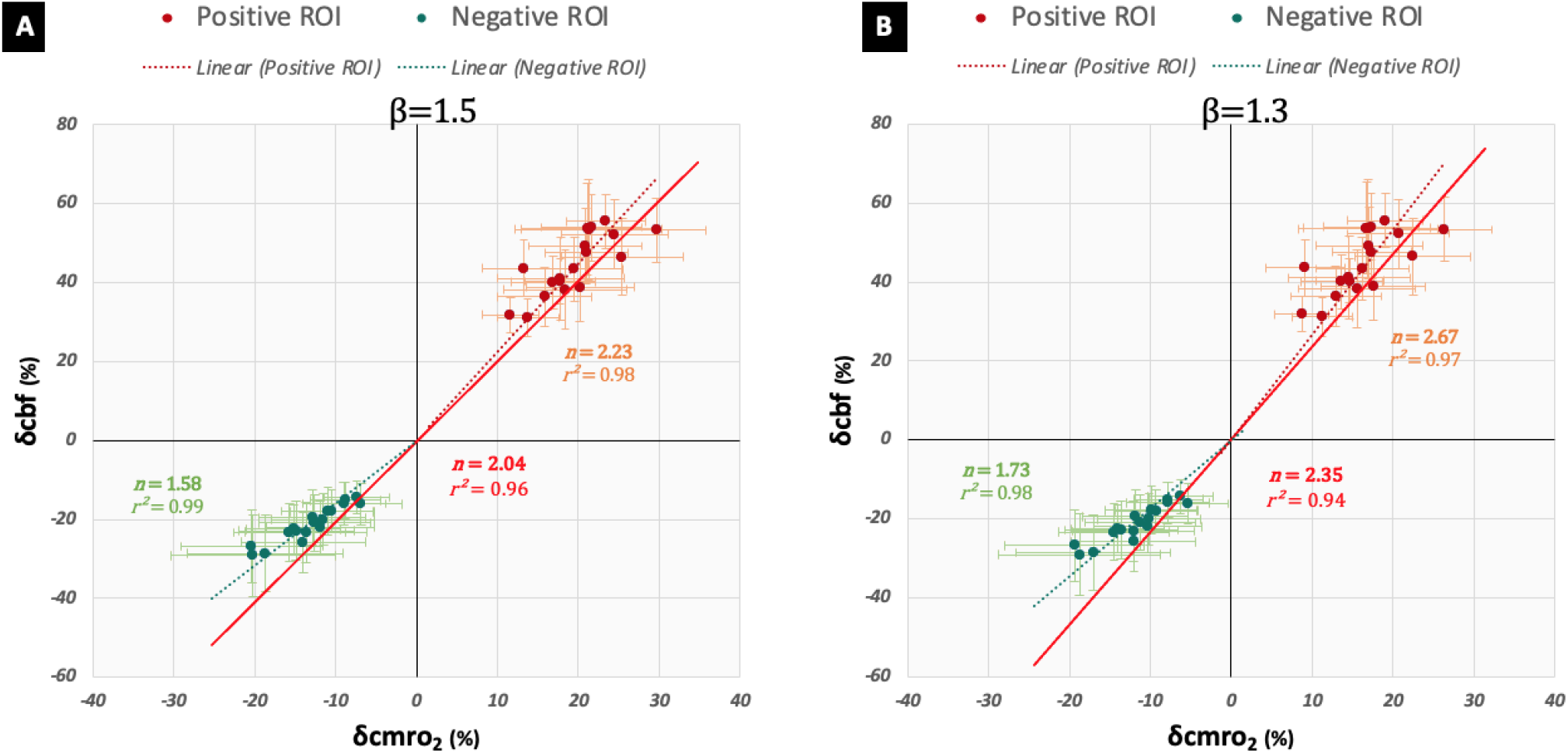
Linear model of flow-metabolism coupling. Plots of δcmro_2_ versus δcbf over both ROIs for δcmro_2_ estimated with *β*=1.5 (A) and *β* = 1.3 (B) for all 18 subjects. The red line indicates the linear fit over all the values, and the dotted lines represent regression lines for values of individual ROIs. Vertical error bars indicate the standard errors (across the ROI) of δcbf and the horizontal error bars represent maximum errors of the δcmro_2_ estimation, propagated from standard errors of δS_BOLD_ and δcbf.

Figures 13, 14 and 15 show implementations of the simplified Wilson Cowan model and its derived models for an increasing *P* with a constant and varying *Q*. The plot of the neuronal population activities for the positive ROI is a replication of Figure 5b in (Buxton, 2021) for *w* = 3. A constant *Q*, however, does not appear to a meaningful description for the negative ROI or deactivation (Figure 13A). With an almost zero inhibitory activity, it translates to a flow-metabolism model where δcmro_2|WC_ ≈ δcbf_WC_ (Figure 14A, red dotted line). Interestingly, no modification of the *Q* value changed this flow-metabolism coupling ratio for the negative values, which is quite improbable as an almost equal change in CBF and CMRO_2_ would never lead to the NBR itself. And unsurprisingly, we see a shift from negative to positive BOLD with decreasing δcbf_WC_ for the Wilson-Cowan δ*S*_BOLD_/δcbf model in Figure 14B. A varying *Q*, on the other hand, appears to work very well for the deactivation model (Figure 13B), resulting in derived models that fit rather well to our estimated and experimental data from the negative ROI (Figures 14 and 15). The corresponding *E*/*I* vs. *P* plots for a linearly increasing *Q* for the positive ROI are not shown. In such cases, the excitatory activity is diminished at high *P* and *Q*, approaching the baseline, while the inhibitory activity only increases. Based on the scaling used in Buxton’s conceptual analysis (Buxton, 2021), equating this *E* and *I* to the physiological parameters will lead to impossibly large values of δcbf (>1,000%). The best solution hence appears to be a model where the *Q* is kept at a constant low value of 0.2 for the activation and one with a varying *Q* for deactivation. The deactivation model was also tested for a positive leak. The *E*/*I* vs. *P* plot for this positive leak (not shown) led to meaningless low minimum excitatory and inhibitory activities (≈10^−165^). Interestingly, once scaled it led to δcmro_2_and δcbf values in the expected ranges, albeit, leading to a slightly higher *n* value than with a negative leakage (Figure 14, green dotted lines). It is to be noted that δcbf_WC_ in the derived models for *β* = 1.3 (Figure 14) and *β* = 1.5 (Figure 15) was calculated with a common 𝒳 = 1.5. This value appears to be a good fit to our data, especially for positive values with *β* = 1.3 and negative values for *β* = 1.5. The model corresponding to a qualitatively better fit for the complementary set of data with 𝒳_NBR_ = 1.1 and 𝒳_PBR_ = 3.2 is highlighted with dashed blue lines in the respective quadrants in Figures 14 and 15.

**Figure 13.**
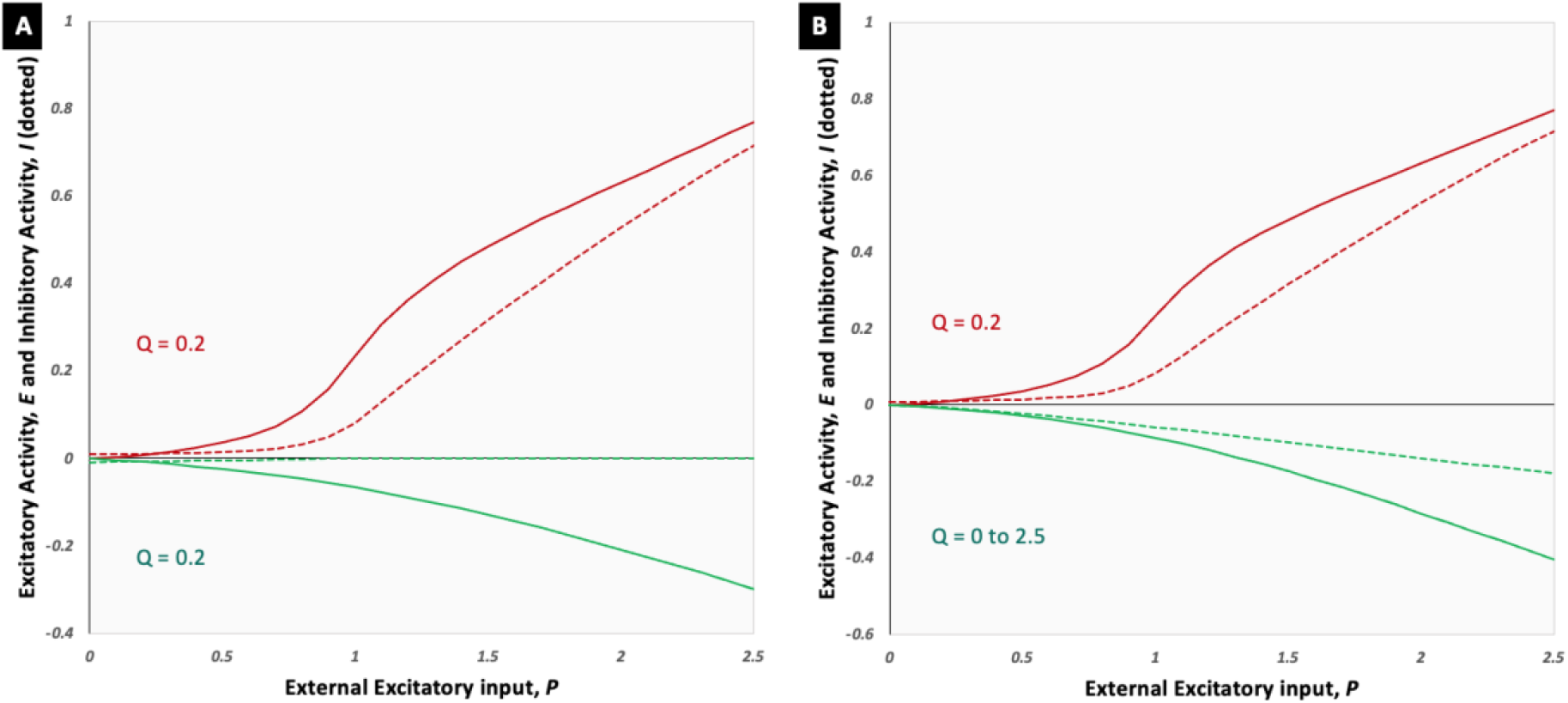
Plot of neuronal population activity against increasing external excitatory input for PBR (red) and NBR (green): (A) constant external inhibitory input Q for both PBR and NBR (B) constant Q for PBR and varying Q for NBR. The bold lines represent excitatory population activity E and the dotted lines represent the corresponding inhibitory activity I.

**Figure 14.**
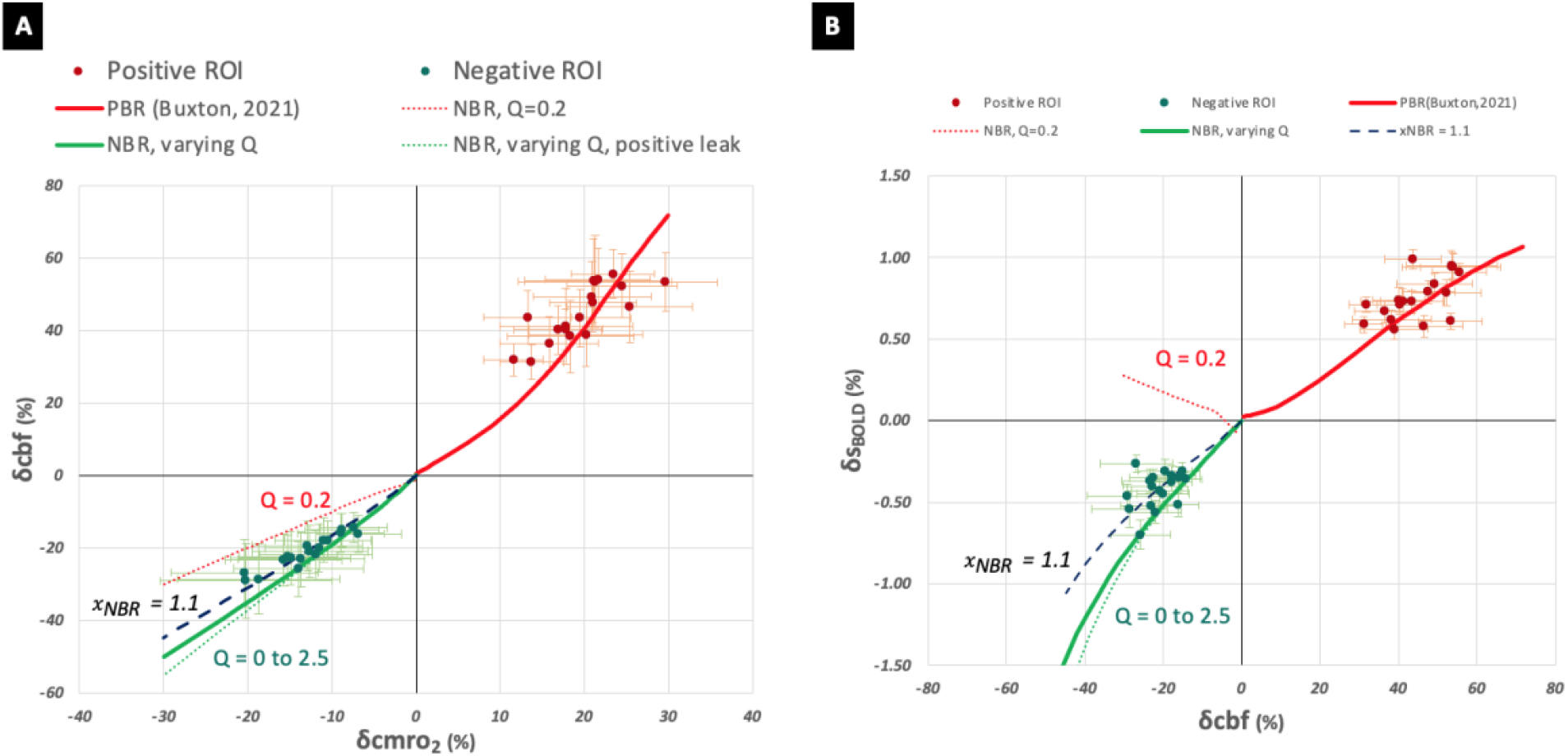
Wilson-Cowan model derived models for *β* = 1.5: (A) flow-metabolism coupling, and (b) the corresponding δS_BOLD_/δcbf. The estimated δcmro_2_ and measured δS_BOLD_ and δcbf in all 18 subjects are represented by red (positive ROI) and green (negative ROI) filled circles. The red lines represent the respective Wilson-Cowan model implemented using a constant Q, while the green lines indicate an implementation for the negative ROI using a linearly varying Q, in bold for a negative leak and dotted for a positive leak. Additionally, the dashed blue lines represent a model with 𝒳_NBR_ = 1.1, all other models shown here used a common 𝒳 value of 1.5. Error bars indicate the standard errors (across the ROI) of δcbf and δS_BOLD_, while horizontal error bars in (a) indicate represent maximum errors of the δcmro_2_ estimation, propagated from standard errors of δS_BOLD_ and δcbf.

**Figure 15.**
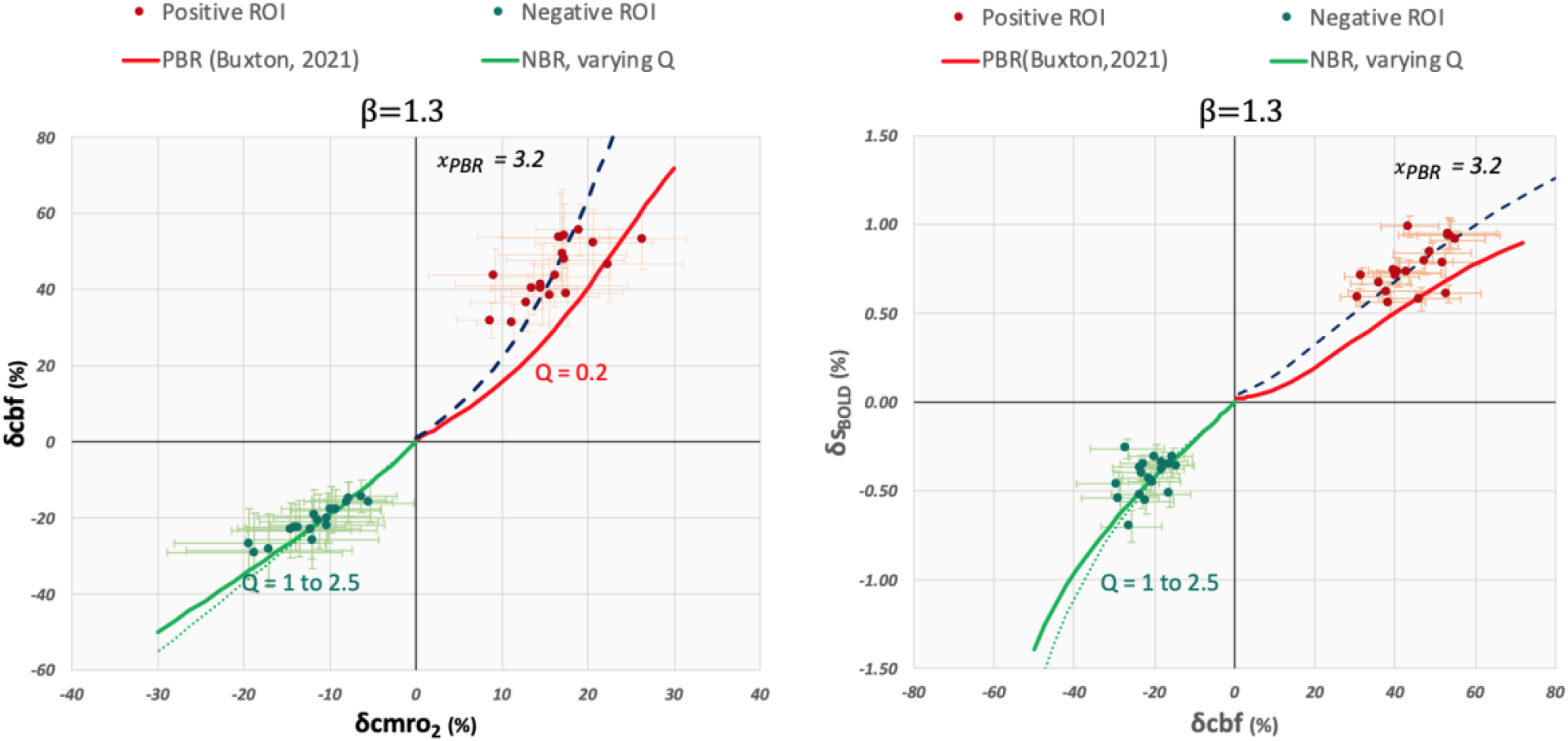
Wilson-Cowan model derived models for *β* = 1.3: (a) flow-metabolism coupling, and (b) the corresponding δS_BOLD_/δcbf. The estimated δcmro_2_ and measured δS_BOLD_ and δcbf in all 18 subjects are represented by red (positive ROI) and green (negative ROI) filled circles. The red lines represent the respective Wilson-Cowan model implemented using a constant Q, while the green lines indicate an implementation for the negative ROI using a linearly varying Q, in bold for a negative leak and dotted for a positive leak. Additionally, the dashed blue lines represent a model with 𝒳_PBR_ = 3._2_, all other models shown here used a common 𝒳 value of 1.5. Error bars indicate the standard errors (across the ROI) of δcbf and δS_BOLD_, while horizontal error bars in (a) indicate represent maximum errors of the δcmro_2_ estimation, propagated from standard errors of δS_BOLD_ and δcbf.

## 4 Discussion

### 4.1 ME-DEPICTING for Simultaneous BOLD- and CBF-fMRI

The DEPICTING readout was recently found to improve the sensitivity in CBF-fMRI when compared to EPI (Devi et al., 2019). In the present study, this superior performance was combined with the capability of simultaneous BOLD-fMRI by the introduction of ME-DEPICTING. A comprehensive study comparing ME versions of both readouts was, hence, conducted, wherein their temporal stabilities and sensitivities for detecting CBF and BOLD responses to an established visual paradigm were investigated.

The use of ME-DEPICTING yielded gains of up to 30% in the functional CNR (Figure 6B) and 20% in the tSNR (Figure 6C) of the ASL contrast compared to ME-EPI. This resulted in substantially larger numbers of supra-threshold voxels for ΔCBF in the first-level analysis (Figure 4). These findings hint on a substantial reduction of unwanted BOLD interference at TE_1_=1.7 ms. Previous work has shown that BOLD contamination is a major deterrent to the detection of functional ΔCBF (Aguirre et al., 2002; Liu and Wong, 2005; Lu et al., 2006; Mildner et al., 2005). Despite the shorter TEs (Figure 2), the *S*_sum_ contrast of ME-DEPICTING provided sufficient sensitivity for simultaneous BOLD measurements, yielding equivalent results as ME-EPI. Therefore, no relevant drop in the BOLD sensitivity was found at the subject level (Figure 4). Interestingly, ME-DEPICTING also yielded more consistently activated regions across subjects for both ΔCBF and BOLD contrast in the analysis of inter-subject variability in Figure 5.

The benefit of shorter TE for the CBF-fMRI, thus, exceeds a potential BOLD sensitivity loss. This can be discussed in terms of differing TE dependencies of both contrasts. The CBF-CNR is linked to the tSNR of the ASL difference signal, which declines with longer TE. The BOLD-CNR, however, is proportional to TE × tSNR (Poser et al., 2006). Because physiological noise exhibits a similar TE dependency as task-induced BOLD signals (Petridou et al., 2009), the SNR gain at longer TE is only moderate, which may explain the comparable BOLD sensitivity of ME-DEPICTING despite shorter TEs.

We, nevertheless, acknowledge that larger, more optimal TE_2_ and TE_3_, equivalent to the ones employed for ME-EPI, could improve the BOLD CNR of the ME-DEPICTING data. This might, however, be not as straightforward. For one, with the recent suggestion of the role of k-space trajectories on BOLD sensitivity and specificity (Engel et al., 2022), there is a possibility that the optimal TEs differ between the two sequences. And more importantly, the temporal efficiency of the DEPICTING sequence suffers for TE > TE_min_. In center-out trajectories, such as DEPICTING, the TE filling is applied twice, and the TE filling delay is not used for data acquisition as with traditional EPI. Achieving the same TE_2_ and TE_3_ as in ME-EPI may, hence, lead to a ‘TR-penalty’ for DEPICTING.

The specifications of the two ME sequences used in the current study were accordingly optimized to allow for meaningful comparisons. For instance, care was taken to obtain an approximately equal TR (ΔTR = 552 ms) while ensuring the lowest possible TE_min_ for both sequences. This was achieved in ME-EPI by the application of partial Fourier-sampling at the cost of somewhat blurrier images (Supplementary Figure S2). We note that in the coarse resolutions used in the present study, such blurring effects are not expected to be of much consequence. The resulting shorter TR_min_ of the ME-EPI (3,182 ms) was then compensated by adjusting the corresponding ‘delay in TR’ to match the TR_min_ of ME-DEPICTING (3,552 ms).

A recommended (Alsop et al., 2015) approach for improving the SNR in ASL is the concept of background suppression (Ye et al., 2000). The suppression of the static-tissue signal is expected to reduce impairments in the ASL difference signal due to physiological and motion-induced fluctuations. The application of such techniques to 2D-ME acquisitions, however, raises two concerns. Firstly, with a single nulling point, it is not possible to obtain a homogenous background suppression across all slices, making it more effective only in combination with single-shot 3D readout techniques (Alsop et al., 2015). Secondly, a reduction in the sensitivity of the simultaneously acquired BOLD signal is another concern. Nevertheless, background suppression has been applied to map CBF and BOLD responses in 2D dual echo (Ghariq et al., 2014) as well as double acquisition (Wesolowski et al., 2009) approaches with reported improvement in CBF-CNR. An appropriate adjustment of the background suppression level was found to keep the reduction in BOLD-CNR within an acceptable range in the former while the BOLD-CNR was preserved in the latter by virtue of a separate acquisition. Recently, a double acquisition sequence with background-suppressed 3D GRASE readout interleaved with a 2D EPI readout allowed for simultaneous measurement of CBF and BOLD, respectively, with a 3-fold increase in ASL tSNR (Fernández-Seara et al., 2016). In our ME-DEPICTING experiments performed without background suppression, an improved CBF -CNR with preserved BOLD sensitivity was achieved by a substantial reduction of TE_1_ and ΔTE.

Besides acquisition-based strategies, retrospective denoising techniques such as ME independent component analysis (ICA) (Kundu et al., 2012) were recently found to improve the functional sensitivities of simultaneously acquired BOLD and CBF contrasts obtained from multi-band ME-EPI data (Cohen et al., 2018). Following an automatic differentiation into BOLD and non-BOLD components by ME-ICA based on their TE dependence, the resulting BOLD and artefactual components are filtered out from TE_1_ and the non-BOLD components from *S*_sum_ data, prior to their statistical analysis. Such data-driven approaches are of particular interest as being unrestricted to a pre-defined model, they could be more tolerable to regional and inter-subject variations. However, due to the stochastic nature of this process, it might compromise a fair sequence comparison. Additional denoising was, thus, omitted, and the data quality was assessed by applying an ASL-specific GLM (Hernandez-Garcia et al., 2010; Mumford et al., 2006), which was capable of regressing out task-specific BOLD signals irrespective of the strength of BOLD fluctuations present in the data.

### 4.2 Relation between Positive and Negative CBF and BOLD Responses

The high sensitivity to CBF changes of ME-DEPICTING along with its capability for a simultaneous BOLD measurement was successfully applied to study the NBR and surrounding regions of a PBR evoked by an established visual stimulus. A difference to previous studies is the use of absolute quantitative measures, such as 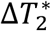, which is more directly related to the deoxyhemoglobin concentration than the BOLD signal itself (Hoge et al., 1999). Additionally, the substantially reduced TE_1_ achieves more accurate measurements of ΔCBF, whereas echo times between 8.2 and 28 ms used in earlier studies (Fukunaga et al., 2008; Mullinger et al., 2014; Stefanovic et al., 2005, 2004; Wilson et al., 2019) are too long to avoid contamination of the ASL signal by the opposing BOLD signal (Aguirre et al., 2002; Liu and Wong, 2005; Lu et al., 2006; Mildner et al., 2005).

The average positive-to-negative BOLD-signal and 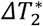 ratio of approximately 2, which agrees very well with other visual activation studies (Pfeuffer et al., 2004; Shmuel et al., 2002, 2001), was also obtained for ΔCBF. This translated to a ratio of ≈1.5 for δcmro_2_ despite using the same assumptions, such as a common *M* value of 4% for both the positive and negative ROIs and for all subjects. In earlier visual PBR studies, absolute ΔCBF values were a more robust measure under varying normal (Kastrup et al., 1999b) or elevated (Li et al., 2000; Whittaker et al., 2016) baseline CBF conditions. In this study, ΔCBF in the negative ROI was found to be independent of CBF_rest_, which could support an extension of the so-called additive hypothesis (Hoge et al., 1999; Li et al., 2000; Sicard and Duong, 2005) to regions of NBR. It implies a constant decrease in ΔCBF in response to stimuli, independent of physiological factors influencing the global CBF_rest_, which if present, would only combine additively. A replication of this independence was, however, not achieved for ΔCBF in the positive ROI. Notably, a significant (*p*=0.01, *r*_2_=0.61) near linear increase of ΔCBF for CBF_rest_ <50 ml/100g/min was revealed, resulting in an overall dependence of both positive ΔCBF and δcbf on CBF_rest_. A potential reason for this effect could be inter-subject ATT variation. Considering the relatively short PLD of 1,200 ms used to obtain an acceptable temporal resolution, we cannot exclude the possibility that only a fraction of the labeled blood water reached the capillaries at long ATTs leading to artificially reduced CBF_rest_ and ΔCBF. The contribution of gender differences to the inter-subject variation cannot be ignored as eight of the nine subjects with CBF_rest_ <50 ml/100g/min in the positive ROI were male. Besides a higher CBF_rest_ (Kastrup et al., 1999a; Rodriguez et al., 1988), ATTs were reported to be shorter by roughly 30% in women compared to men (Liu et al., 2012). The invariance in the negative ROI would be in line with this interpretation, because the ATT increases from V1 towards the border zone of the posterior perfusion territory in extrastriate regions (most of the positive ROI) (Mildner et al., 2014). However, considering a cortical thickness of about 2 mm in human V1 (Alvarez et al., 2019), a residual bias of the CBF measurements by partial-volume effects causing artificially reduced ΔCBF and CBF_rest_ values can also not be excluded. Additionally, as transit effects do not impact 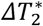, there was no variation with CBF in both ROIs. 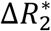 would hence, combine additively.

In the quantitative comparison of the positive and negative responses, both ROIs were carefully kept as homogenous as possible by the application of a 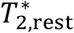 mask, which is expected to reduce differences due to the presence of pial veins and CSF. Venous contributions were further kept to a minimum by only selecting ROIs with concurrently significant activation or deactivation with both BOLD- and CBF-fMRI. The differences in 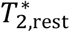 and CBF, however, highlights the structural differences between the two ROIs, as the negative ROI was located entirely in V1, which is known to have a higher vascular density (in particular, in layer IV) than extrastriate visual areas, which make up the positive ROI together with foveal V1 (Weber et al., 2008). It is suggested in Figure 9B that the observed difference in the average coupling ratio δ*S*_BOLD_/δcbf between the positive and negative ROI is caused by their differing mean CBF_rest_. The higher average δ*S*_BOLD_/δcbf in the negative ROI, which is in line with an earlier NBR study in V1 (Shmuel et al., 2002), is, thereby, explained by a higher CBF_rest_ in peripheral V1. This demonstrates the benefit of CBF quantification for a consistent interpretation of combined CBF and BOLD studies. It further suggests that coupling ratios derived from relative CBF changes cannot easily be compared between brain regions (Whittaker et al., 2016).

The ΔCBF timecourses closely followed their 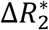 counterparts, albeit with less pronounced post-stimulus transients. Typical BOLD temporal characteristics were seen in the 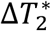 timecourses (Figures 10 and 11) with the negative 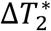 transients bearing a striking resemblance to previously published NBR transients in humans (Huber et al., 2014) and non-human primates (Shmuel et al., 2006) at higher field strengths. Despite the low temporal resolution (approx. 7s), the elusive PSU and PSO of the, respectively, positive and negative ΔCBF responses were detected in most subjects. The general shape of the positive ΔCBF transient with a subtle PSU (*n* =8), which was occasionally (*n* =3) followed by another overshoot, agrees reasonably well with transients reported in recent literature (Kim et al., 2020; Mullinger et al., 2017, 2014, 2013). The TTP_1_ of the positive BOLD and ΔCBF responses were also similar in these studies. Remarkably, the shape of the signals obtained here using a stimulation period of approximately 28 s did not differ much from previously published ones obtained with a brief 2s visual stimulation (Kim et al., 2020). While a tight CBF-BOLD coupling is expected in fast fMRI dynamics, the strong correlation of the two signals in our long stimulation study could be construed to their microvasculature origins (Polimeni and Lewis, 2021). It is possible that, despite our rather low resolution, the utilization of a common overlap along with the additional constraints placed to correct for partial voluming effects rendered the ROIs microvasculature dominant.

Differences in the shapes of the PBR and NBR have been widely reported (Klingner et al., 2015). The combination of pCASL with a sensitive short-TE readout allowed to demonstrate the mirroring of this distinction in the ΔCBF signals. While this finding once again verifies the role of vasodilation and vasoconstriction in the PBR and NBR, respectively, it also provides a strong indication to the inhibitory origin of the NBR and the corresponding ΔCBF signals measured here. NBR temporal dynamics similar to that of the negative 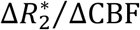 signals had previously been found to correlate strongly with simultaneously measured inhibitory neural responses in monkey visual cortex (Shmuel et al., 2006). The strong correspondence of the negative ΔCBF signal to 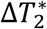 then, by extension, suggests an inhibitory control to the reduced blood flow. Additionally, the faster TTP of the negative signals is in agreement with recent findings of an earlier peak of the inhibitory hemodynamic response function (HRF) than that of the excitatory HRF observed in genetically modified mice at 15.7 T (Moon et al., 2021). Faster inhibitory neurovascular dynamics had earlier been predicted to explain the faster evolution of the NBR signals (Havlicek et al., 2017). The disparity in the shape of the positive and negative 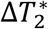 signals can also be attributed to their differing spatial origins. Recent high-resolution studies in macaques (Goense et al., 2012) and humans (Huber et al., 2014) have reported differing laminar origins of the PBR and the NBR, with the former peaking at the cortical surface and the latter at the deeper layers. HRFs at the deeper layers and hence away from pial vasculature had been found to have shorter amplitudes and faster TTP^1^ than at the superficial layers for the positive response (Siero et al., 2011). The exhibition of similar dynamics in the measured negative signals, hence, suggests a larger concentration of inhibitory interneurons along with a faster vasoconstriction of the microvasculature dominant deeper layers. The higher specificity would explain the stronger correlation between NBR and the neuronal response, the higher ratio of BOLD to neural response amplitudes (Shmuel et al., 2006) and the determination of a larger neuronal contribution to the overshoot of the NBR (Havlicek et al., 2017), contrary to that of the PBR. This is also reflected in the significantly differing coupling ratios of the mean amplitudes (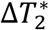 and δ*S* /δcbf) between the positive and negative ROIs. Despite the strong correspondence in the shape of the 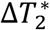 and ΔCBF in their individual domain, decreases in CBF appear to warrant a larger change in NBR than CBF increases in PBR. These differences in spatio-temporal excitatory and inhibitory hemodynamics with regards to CBF reinforces previous evidence of a differing neurovascular coupling between PBR and NBR (Huber et al., 2014; Mullinger et al., 2014).

The significantly differing *n* (δcbf/δcmro_2_) values between the two ROIs, irrespective of the *β* value, further supports this train of thought (Figure 12). This is in agreement with recent literature, wherein, the difference persisted with *n*_NBR_ being consistently lower than *n*_PBR_ regardless of the *M* value or model parameters (Mullinger et al., 2014). But it should be noted that the different scaling behaviour of the δ*S*_BOLD_ -δcbf grid onto the δcmro_2_ -δcbf plane (Chiarelli et al., 2007) in case of activation and deactivation introduces a larger bias in the derived δcmro_2_ values which might also contribute to the observed differences in *n*. We further note that due to the many assumptions made, the application of the Davis model to our data serves only as a means for qualitative assessment. Having the same positive and negative ROI would, however, avoid the difficulties of having different CBF_rest_ and possibly also *M* values (Ances et al., 2007; Chiarelli et al., 2007).

The replication of the Wilson-Cowan model for flow-metabolism coupling (Buxton, 2021) and its adaptation to regions of NBR, however, provides new insights into this matter. On account of the large errors, the original assumption of 𝒳_PBR_ = 1.5 appeared to fit rather well to our multi-subject normocapnic data in the positive ROI for *β* = 1.5 (Figure 14). The apparent failure of a corresponding deactivation model with a constant *Q* (Figure 14), suggests a difference in the neuronal contribution between PBR and NBR. As hypothesized by others (Mullinger et al., 2014), it seems plausible that other inhibitory mechanisms are relevant in the NBR that are not present in the PBR. Different 𝒳 values also appear to qualitatively fit the experimental data better than a common value for both ROIs (Figures 14 & 15). This might hint at a differing neuronal control of CMRO_2_ and CBF changes in the NBR. Furthermore, the substantial impact of the choice of the *β* value used for the δcmro_2_ estimation is highlighted, as this fit was clearly sub-optimal for the PBR data estimated with *β*=1.3 (Figure 15A). However, based on the number of assumptions we have made, such an inference can only be taken with more than a pinch of salt. The lack of data in the lower range and higher range for the positive and negative ROI, respectively, and the disparate anatomical locations of the two ROIs are also limiting factors. It is, hence, very important to note here that these are just mere conjectures. The other assumptions used for the positive ROI in the model might not work for the negative ROI, namely, the gain or threshold of the sigmoid function, identical value of the weights or the omission of a self-inhibitory input. Moreover, errors in the quantitative estimations cannot be overlooked. Nevertheless, such models provide an intriguing perspective and could complement our understanding of neuronal dynamics relating to the CMRO_2_ and CBF changes accompanying the BOLD response, both positive and negative.

### 4.3 Limitations

The anatomically differing positive and negative ROIs poses a major constraint in the interpretation of results presented in the current study. Their disparity in basal physiology was reflected in the significantly higher CBF_rest_ obtained for the negative ROIs. Apart from the CBF_rest_ dependence of δcbf and, consequently, potential effects on the δs_BOLD_/δcbf coupling ratio, baseline CBF levels are also known to influence the temporal dynamics of the BOLD signal (Behzadi and Liu, 2005; Cohen et al., 2002; Kemna and Posse, 2001). An obvious solution to avoid these regional variations would be to measure PBR and NBR from the same ROI. Such investigations would then rely on combinations of stimuli (as in Shmuel et al., 2006) that would evoke these responses through their sequential presentation, ideally within the same run. Care, however, should be taken when selecting the appropriate inter-stimulus intervals (ISI) in these paradigms.

It is to be noted that the CBF_rest_ in the present study were estimated from the resting blocks within the paradigm. Although these blocks were longer (1.5×) than the task blocks to allow approximation to a baseline state, it is possible that the presence of oscillatory post-stimulus transients made these ISIs insufficient. A comparison of these CBF_rest_ values with the ones measured from the corresponding resting-state runs (*substudy 1*) in subjects P6, P8 and P9 over both the positive and negative ROIs proved insignificant (*p*=0.62). CBF_rest_ over the negative ROI was also found to be higher than that in the positive ROI, with the exception of P6, wherein with a negative ROI comprising of only 21 voxels, the CBF_rest_ over the positive ROI was slightly higher (Supplementary Table S3). A prolonged resting period within the same run, at the start or end of the functional paradigm is expected to provide a better estimate of the baseline condition.

Partial overlaps between regions of CBF increases and PBR are well reported (Stefanovic et al., 2004). This difference in the spatial extent of activation have been attributed to the higher SNR of the BOLD signal and the varying sources of contrast of the two signals, with BOLD arising from a more venous source contrary to the capillary tissue bed from which CBF changes are measured (Lipton et al., 2000; Luh et al., 2000). The above hypotheses could apply to the NBR data and the corresponding decreases in CBF. The negative ROIs generated here had an average spatial overlap of ∼51% compared to ∼90% of the positive ROIs and could, hence, be explained by the inherently lower sensitivities of the two negative signals. Apart from the lower amplitudes, the larger mismatch between the canonical HRF employed in the GLM and the shape of the evoked negative signals (Figure 11) could have attributed to their lower SNR and larger variance (Greve et al., 2013; Mullinger et al., 2014). The larger reduction in spatial overlap might also result from a prolonged ATT brought about by the reduced blood flow rendering the PLD of 1,200 ms insufficient, causing the detection of significant negative ΔCBF from arteries rather than brain tissue. Superposition with an arterial probability map (Mouches and Forkert, 2019) over regions of significant ΔCBF decreases that did not overlap with the NBR indicated this to be the case for some of the voxels. A follow-up study employing two ASL acquisitions with different PLDs would clarify the potential influence of this effect. pCASL scans with multiple PLDs for each subject would have also helped ascertain inter-subject difference in ATT. However, as these scans are very time-consuming and the full extent of the gender differences not initially comprehended, they were omitted from the present study. Subsequent experiments would likely involve such a scan in a prior session. ATT-insensitive approaches such as VS-ASL (Wong et al., 2006) and the more recently introduced VESPA ASL (Woods et al., 2022) which additionally provides measures of ATT are interesting alternatives to consider in this regard. The study could have also benefitted from retinotopic scans which would have substantiated the anatomical locations of the positive and negative ROIs.

The extent of spatial overlap is also expected to vary with the statistical thresholds applied. The application of a statistical threshold filters out insignificant voxels, but it also biases average values in a thereby defined ROI by introducing an arbitrary minimum effect size. It is, hence, important to note that although the size of the positive and negative ROIs changed, all correlations reported in the current study were reproduced when the applied statistical threshold was altered.

## 5 Conclusion

A pCASL-prepared ME-DEPICTING sequence achieving short TE_1_ and ΔTE was introduced. In comparison to standard ME-EPI, an improved sensitivity for mapping functional CBF changes was obtained without a relevant CNR drop of the simultaneously acquired BOLD response. The method was employed at 3 T for a detailed investigation of negative CBF and BOLD responses in peripheral V1 and their comparison with simultaneously evoked positive responses in foveal V1. Significant differences between the two ROIs were revealed in terms of their coupling ratios δ*S*_BOLD_/δcbf as well as the corresponding flow-metabolism coupling factor *n*. However, an influence of different CBF_rest_ levels found in both regions on these ratios was also shown, the extent of which requires further investigation. Differences were further detected in the shapes of the positive and negative transients when comparing their primary and post-stimulus peaks. Lastly, the metabolic load and contributing neuronal dynamics of the two ROIs were assessed to gain a better understanding into the underlying physiology.

## Data and code availability statement

Pre-processed raw data, their derived data from multi-echo fitting, and design matrices used for the analyses will be made available upon acceptance at https://osf.io/3s89p/. The pre-processing and analyses steps have been well documented in the Methods section and were mostly executed using publicly available Statistical Parametric Mapping (SPM) (https://www.fil.ion.ucl.ac.uk/spm/software/spm12/) and FMRIB Software Library (FSL)(http://www.fmrib.ox.ac.uk/fsl) utilities. All data required for CBF quantification have also been provided. If required, the IDL scripts used for this purpose can be made available upon request.

## CRediT authorship contribution statement

**Ratnamanjuri Devi:** Conceptualization, Methodology, Software, Formal analysis, Investigation, Writing - original draft, Writing - review & editing, Visualization. **Jöran Lepsien:** Methodology, Software. **Kathrin Lorenz:** Methodology. **Torsten Schlumm:** Methodology, Software, Data curation. **Toralf Mildner:** Conceptualization, Methodology, Software, Investigation, Writing -original draft, Writing - review & editing. **Harald E. Möller:** Conceptualization, Investigation, Writing - original draft, Writing - review & editing, Supervision, Project administration, Funding acquisition.

## Acknowledgement

We would like to thank Dr. Laurentius Huber for providing us with his extensive notes on the stimulus used in the current study and Dr. Joseph Whittaker for helpful discussions. We also thank Anke Kummer and Simone Wipper for their assistance with the volunteers. This work has been supported by the Max Planck Society and by the International Max Planck Research School on Neuroscience of Communication: Function, Structure, and Plasticity.

