## Supplementary Material for "Multi-Echo Investigations of Positive and Negative CBF and Concomitant BOLD Changes"

### Abbreviations

2D, 3D = two-, three-dimensional; ASL = arterial spin labeling; BET = Brian Extraction Tool; BOLD = blood oxygenation level dependent; CNR = contrast-to-noise ratio; CSF = cerebrospinal fluid; DEPICTING = Double shot EPI with Center-out Trajectories and Intrinsic NaviGation; EPI = echo planar-imaging; fMRI = functional magnetic resonance imaging; FSL = FMRIB Software Library; GLM = general linear model; GM = gray matter; GRAPPA = GeneRalized Autocalibrating Partial Parallel Acquisition; GRE = gradient-recalled echo; HRF = hemodynamic response function; ICE = Image Calculation Environment; ISI = Inter-stimulus Interval; ME = multi echo; MNI = Montreal Neurological Institute; MP-RAGE = Magnetization-Prepared RApid Gradient Echo; MP2RAGE = Magnetization-Prepared 2 Rapid Acquisition Gradient Echoes; NBR = negative BOLD response; PBR = positive BOLD response; pCASL = pseudo-continuous ASL; PLD = post-labeling delay; PSO = post-stimulus overshoot; PSU = post-stimulus undershoot; RF = radiofrequency; ROI = region of interest; SNR = signal-to-noise ratio; tSNR = temporal SNR; $TTP_1$ = time to peak of the primary signal; $TTP_2$ = time to peak of the post-stimulus transient; V1, V2, V3, V4, V5 = visual area V1, V2, V3, V4, V5.

### Mathematical symbols

$a$: gain,

$\Delta B_0$: main magnetic field offset,

$B_{1,\mathrm{av}}$: average RF pulse amplitude,

CBF: cerebral blood flow,

$\mathrm{CBF}_{\mathrm{rest}}$: CBF at rest,

$\mathrm{CBF}_{\mathrm{stim}}$: CBF during stimulation,

ΔCBF: absolute stimulus-induced CBF change,

δcbf: relative stimulus-induced CBF change,

CBV: cerebral blood volume;

$\mathrm{CMRO}_2$: cerebral metabolic rate of oxygen consumption;

$\delta\mathrm{cmro}_2$: relative stimulus-induced $\mathrm{CMRO}_2$ change,

$E$: excitatory activity,

$G_{\text{av}}$: average gradient amplitude,

$I$: inhibitory activity,

$n$: integer; also, flow-metabolism coupling factor,

$P$: external excitatory input,

$p$: error probability,

$\langle p \rangle$: average error probability,

$Q$: external inhibitory input,

$r^2$: coefficient of determination (squared Pearson correlation coefficient),

$R_2^*$: effective transverse relaxation rate,

$R_{2,\text{rest}}^*$: effective transverse relaxation rate at rest,

$R_{2,\text{stim}}^*$: effective transverse relaxation rate during stimulation,

$\Delta R_2^*$: change of the effective transverse relaxation rate between stimulation and rest,

$S_0$: signal intensity at zero echo time,

$S_n$: signal intensity at echo time $\text{TE}_n$,

$S_{\text{rest}}$: signal intensity at rest,

$S_{\text{stim}}$: signal intensity during stimulation,

$S_{\text{sum}}$: signal intensity obtained by weighted summation of multiple echoes,

$\text{StdErr}(x)$: standard error of $x$,

$\delta s_{\text{BOLD}}$: relative BOLD signal change,

$\mathcal{S}(x)$: sigmoidal transfer function in the Wilson-Cowan model,

$T_1$: longitudinal relaxation time,

$T_2^*$: effective transverse relaxation time,

$T_{2,\text{rest}}^*$: effective transverse relaxation time at rest,

$\Delta T_2^*$: change of the effective transverse relaxation time between stimulation and rest,

$T_{\text{acq}}$: acquisition time,

TE: echo time,

$\text{TE}_{\text{eff}}$: effective echo time,

$\text{TE}_n$: echo time of the $n$-th echo,

$\text{TE}_{\text{min}}$: minimum echo time,

$\Delta\text{TE}$: inter-echo time,

TR: repetition time,

$TR_{min}$: minimum repetition time,

$t$: time,

$w$: strength of synaptic connections,

$x_{PBR}, x_{NBR}$: relative weightings of inhibitory activity contributing to changes in CBF (Wilson-Cowan model) in the PBR and NBR, respectively,

$\alpha$: Grubbs exponent relating cerebral blood flow and volume changes (Davis model),

$\beta_i$: GLM parameter,

$\beta$: exponent in the Davis model that varies with field strength and vascularization,

$\tau_0$: decay time constant,

$\theta$: threshold.

### Figures

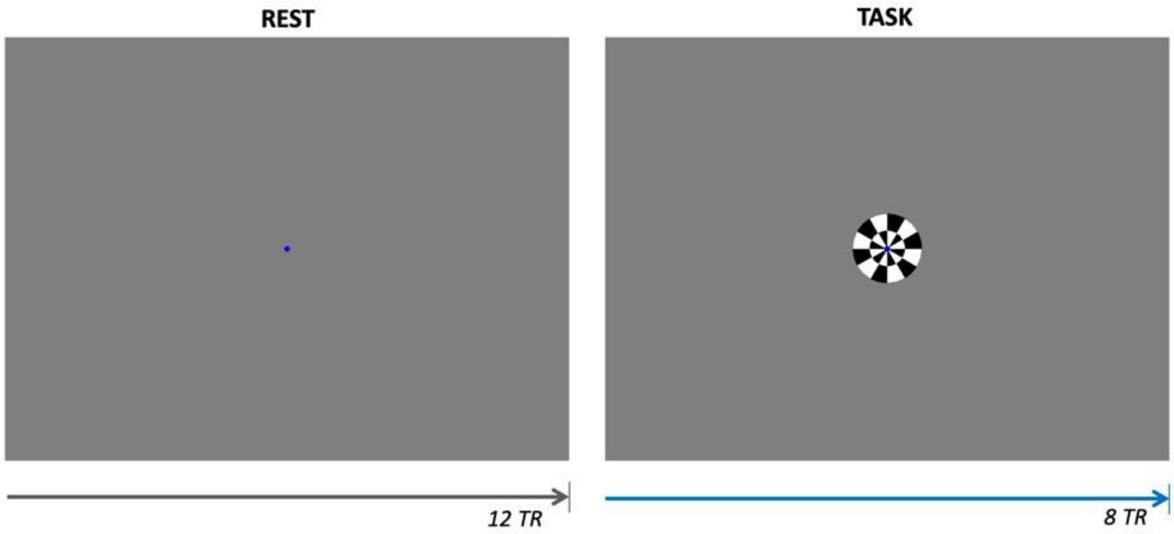


***Supplementary Figure S1.*** *Visual stimulus used to induce a central NBR in the primary visual cortex surrounded by areas of a PBR. A central fixation point was present at all times.*

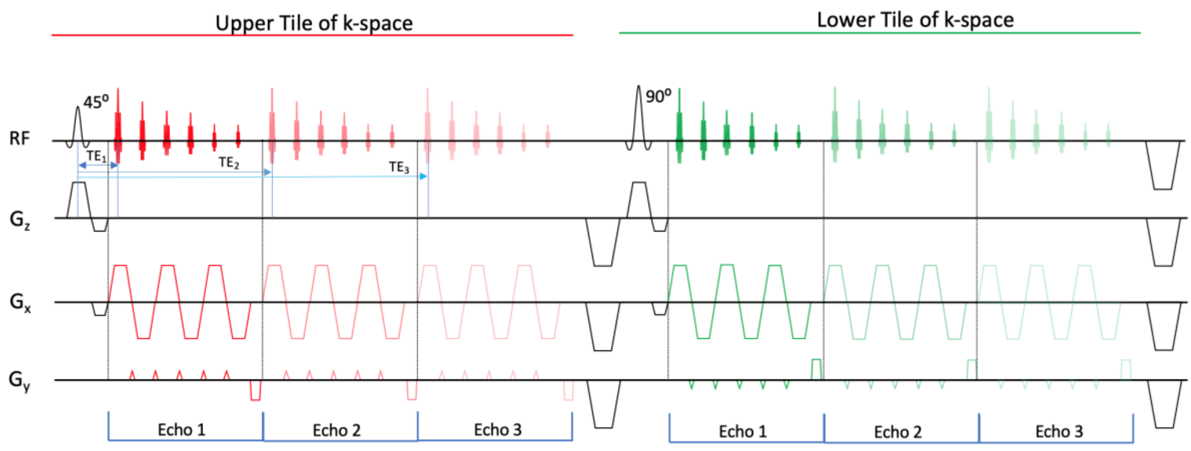


***Supplementary Figure S2.*** *Schematic of the ME-DEPICTING sequence for the acquisition of 3 echoes. Note the lack of acquisition during the TE filling times.*

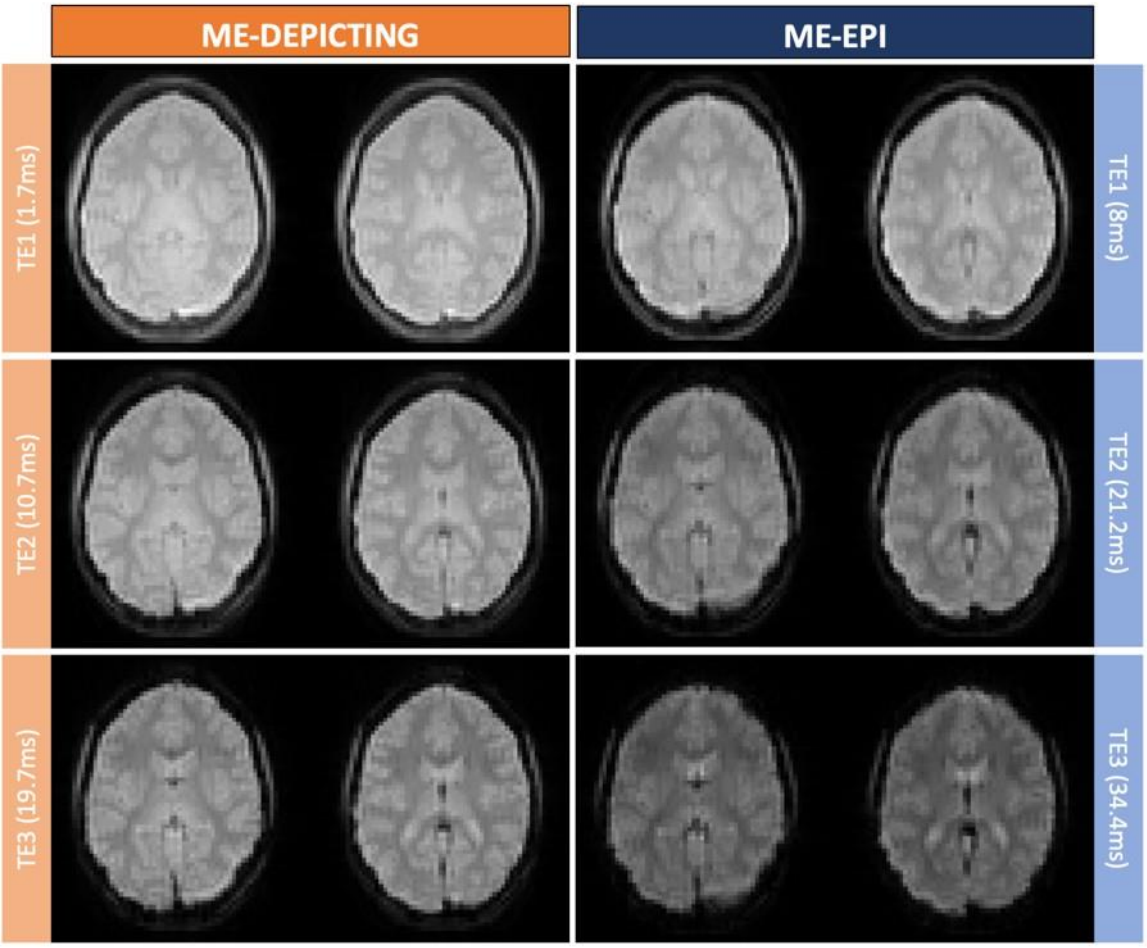

***Supplementary Figure S3.*** *Raw images of the three echoes obtained with ME-DEPICTING **(A)** and ME-EPI **(B)** in the same subject in slices containing the visual cortex.*

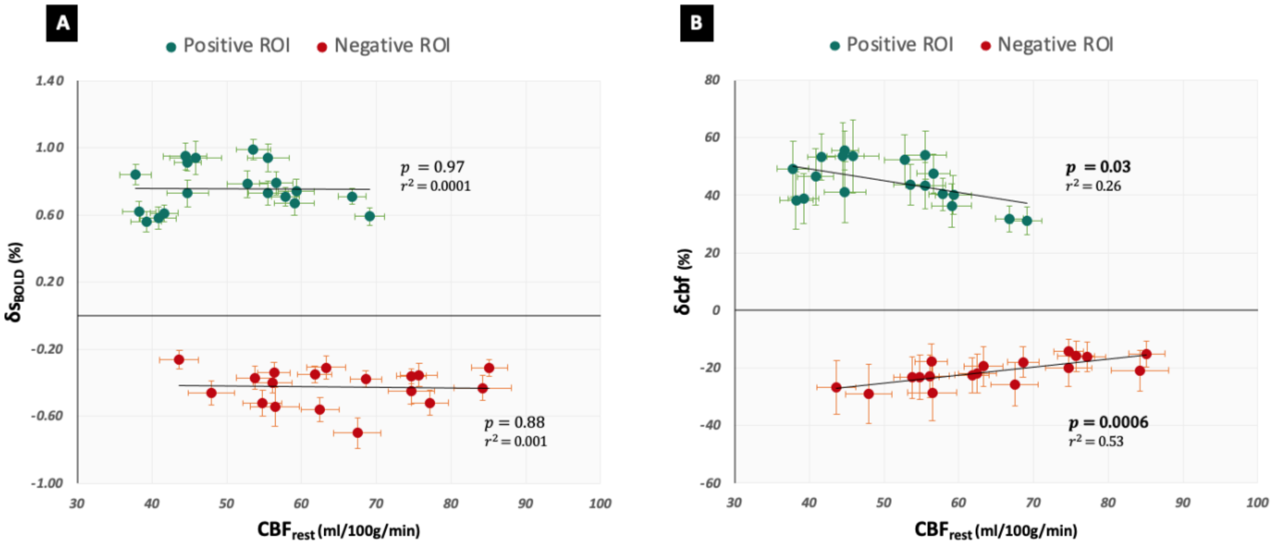


***Supplementary Figure S4.*** *Stimulus-induced $\delta s_{\mathrm{BOLD}}$ **(A)** and $\delta cbf$ **(B)** plotted as a function of their corresponding $CBF_{rest}$ for all subjects. Green and red circles show average values in the positive and negative ROIs, respectively. Error bars indicate the standard errors (across the ROI) of $\delta s_{\mathrm{BOLD}}$, $\delta cbf$ and $CBF_{rest}$.*

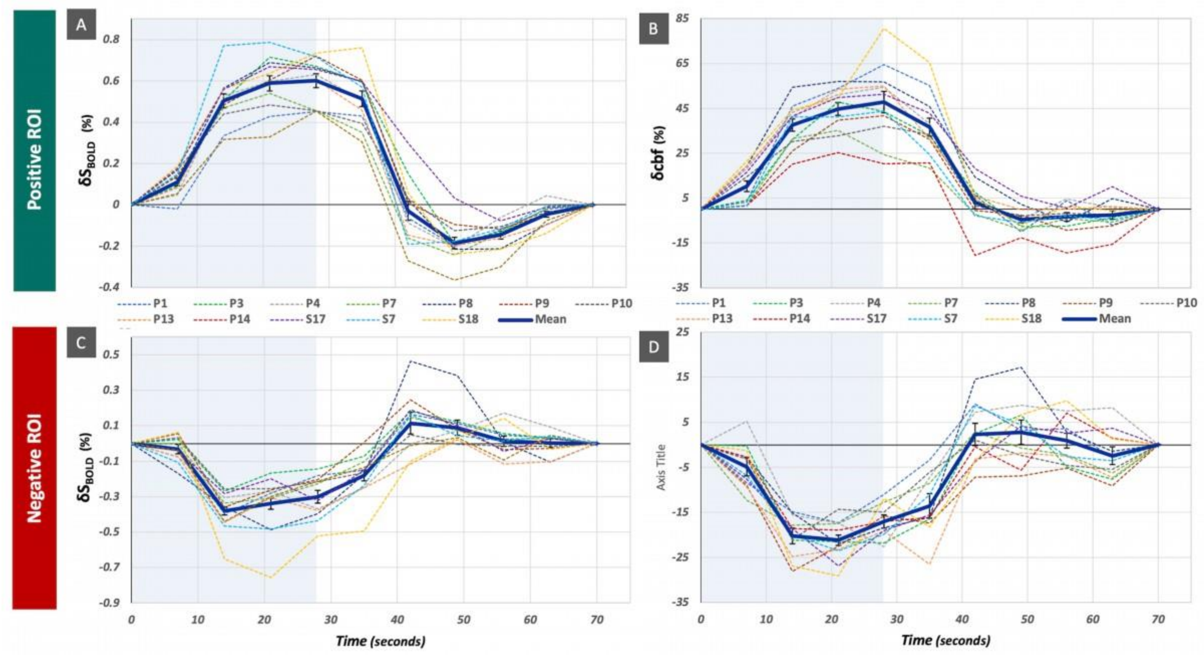


***Supplementary Figure S5.*** *Cycle-averaged $\delta s_{\mathrm{BOLD}}$ **(A, C)** and $\delta cbf$ **(B, D)** timecourses in twelve subjects with a minimum ROI size of 100 voxels in positive **(A, B)** and negative ROIs **(C, D)** ROIs, namely P1, P3, P4, P7, P8, P9, P10, P13, P14, S17, S7 and S18. Subject-averaged mean timecourses are shown as bold solid blue lines. The error bars indicate one standard error of the mean (SEM). Blue shaded regions indicate the duration of the task.*

### Tables

| *Subjects* | *Significantly activated voxels* | | **Positive ROI** (voxels) | **% overlap** | *Significantly deactivated voxels* | | **Negative ROI** (voxels) | **% overlap** |
|---|---|---|---|---|---|---|---|---|
| | **BOLD** | **CBF** | | | **BOLD** | **CBF** | | |
| ***P1*** | 3086 | 2309 | 2087 | 90 | 2717 | 953 | 801 | 84 |
| ***P2*** | 3230 | 1755 | 1661 | 95 | 533 | 107 | 6 | 6 |
| ***P3*** | 4886 | 4099 | 3728 | 91 | 3364 | 1967 | 1000 | 51 |
| ***P4*** | 3803 | 3596 | 3109 | 86 | 1753 | 773 | 235 | 30 |
| ***P5*** | 4581 | 3483 | 3093 | 89 | 896 | 112 | 14 | 13 |
| ***P6*** | 1440 | 1111 | 908 | 82 | 1046 | 70 | 21 | 30 |
| ***P7*** | 5950 | 5077 | 4611 | 91 | 2241 | 972 | 703 | 72 |
| ***P8*** | 3865 | 2594 | 2486 | 96 | 3305 | 302 | 232 | 77 |
| ***P9*** | 4108 | 3348 | 2932 | 88 | 2025 | 698 | 272 | 39 |
| ***P10*** | 3338 | 2821 | 2628 | 93 | 2215 | 760 | 543 | 71 |
| ***P11*** | 2564 | 2138 | 1681 | 79 | 1373 | 199 | 47 | 24 |
| ***P12*** | 3030 | 4228 | 2732 | 65 | 519 | 118 | 72 | 61 |
| ***P13*** | 2880 | 2144 | 1993 | 93 | 2919 | 809 | 365 | 45 |
| ***P14*** | 761 | 384 | 357 | 93 | 1499 | 139 | 101 | 73 |
| ***P15*** | 2643 | 1360 | 1266 | 93 | 1701 | 103 | 64 | 62 |
| ***S16*** | 4611 | 2352 | 2226 | 95 | 503 | 21 | 0 | 0 |
| ***S17*** | 4788 | 3060 | 2715 | 89 | 1592 | 248 | 170 | 69 |
| ***S7*** | 6198 | 3687 | 3590 | 97 | 2665 | 564 | 427 | 76 |
| ***S18*** | 2509 | 1916 | 1843 | 96 | 5437 | 1974 | 1712 | 87 |
| ***Mean ± SD*** | **3593 ± 1393** | **2709 ± 1178** | **2402 ± 1026** | **90 ± 8** | **2016 ± 1224** | **573 ± 591** | **357 ± 440** | **51 ± 27** |

***Supplementary Table S1.*** *Single-subject and group averages plus/minus one standard deviation (SD) of the number of voxels with significant BOLD ($p < 10^{-4}$) and CBF ($p < 0.01$) activation and deactivation, sizes of the respective overlapping regions i.e., Positive and Negative ROIs and the % overlap of this ROI with CBF activation and deactivation.*

| *Subjects* | | *P1* | *P2* | *P3* | *P4* | *P5* | *P6* | *P7* | *P8* | *P9* | *P10* | *P11* | *P12* | *P13* | *P14* | *P15* | *S16* | *S17* | *S7* | *S18* | ***Mean ± SD*** |
|---|---|---|---|---|---|---|---|---|---|---|---|---|---|---|---|---|---|---|---|---|---|
| $\Delta R2^*/\Delta CBF$ $s^{-1}/(ml/100g/min)$ | *Positive ROI* | 0.024 | 0.032 | 0.024 | 0.026 | 0.030 | 0.024 | 0.027 | 0.038 | 0.025 | 0.022 | 0.032 | 0.024 | 0.025 | 0.038 | 0.030 | 0.034 | 0.027 | 0.036 | 0.026 | **0.029 ± 0.005** |
| | *Negative ROI* | 0.025 | 0.014 | 0.018 | 0.02 | 0.020 | 0.015 | 0.026 | 0.029 | 0.025 | 0.019 | 0.024 | 0.018 | 0.022 | 0.025 | 0.021 | - | 0.018 | 0.033 | 0.032 | **0.023 ± 0.005** |
| $\delta s_{BOLD}/\delta cbf$ | *Positive ROI* | 0.011 | 0.014 | 0.018 | 0.017 | 0.016 | 0.012 | 0.022 | 0.017 | 0.018 | 0.019 | 0.018 | 0.015 | 0.017 | 0.016 | 0.018 | 0.018 | 0.018 | 0.023 | 0.017 | **0.017 ± 0.003** |
| | *Negative ROI* | 0.021 | 0.016 | 0.016 | 0.016 | 0.022 | 0.01 | 0.025 | 0.023 | 0.032 | 0.021 | 0.019 | 0.021 | 0.022 | 0.019 | 0.016 | - | 0.018 | 0.026 | 0.027 | **0.020± 0.005** |

***Supplementary Table S2.*** *Single-subject and group averages plus/minus one standard deviation (SD) of $\Delta R_2^*/\Delta CBF$ and $\delta s_{BOLD}/\delta cbf$ ratios over the positive and negative ROIs.*

| Subjects | | Number of voxels | | $CBF_{rest}$ (ml/100g/min) | |
|---|---|---|---|---|---|
| | | *Positive ROI* | *Negative ROI* | *Positive ROI* | *Negative ROI* |
| ***P6*** | *substudy 2* | 908 | 21 | **40.8 ± 12.3** | **43.6 ± 14.0** |
| | *substudy 1 (Resting run)* | 908 | 21 | **51.5 ± 13.8** | **47.0 ± 12.5** |
| ***P8*** | *substudy 2* | 2486 | 232 | **37.8 ± 13.0** | **54.8 ± 13.0** |
| | *substudy 1 (Resting run)* | 2486 | 232 | **35.6 ± 17.1** | **46.4 ± 20.4** |
| ***P9*** | *substudy 2* | 2932 | 272 | **59.3 ± 17.6** | **77.2 ± 10.3** |
| | *substudy 1 (Resting run)* | 2932 | 272 | **62.6 ± 15.4** | **78.6 ± 10.5** |

***Supplementary Table S3.*** *Comparison of $CBF_{rest}$ (mean ± SD) obtained from the positive and negative ROIs of substudy 2 with corresponding values from resting state runs of substudy1 in three subjects.*

### Methods

The Wilson Cowan model (Wilson and Cowan, 1972) describes a pair of non-linear differential equations that model the dynamics of spatially localized populations containing both excitatory and inhibitory neuronal populations. The excitatory and inhibitory populations are represented in these equations as $E$ and $I$, respectively, with the remaining key parameters being the decay time constant ($\tau_0$); the synaptic weights representing the strength of connectivity between the different subpopulations, self-excitation as well as self-inhibition ($w_{EI}, w_{IE}, w_{EE}, w_{II}$); the synaptic threshold ($\theta$); the gain of the system ($a$); the refractory time constant ($r$), external excitatory ($P$) and inhibitory inputs ($Q$):

$$\tau_0 \frac{dE}{dt} = -E + (1 - rE)\, \mathcal{S}(w_{EE}E - w_{IE}I + P), \quad \text{(S1a)}$$

$$\tau_0 \frac{dI}{dt} = -I + (1 - rI)\, \mathcal{S}(w_{EI}E - w_{II}I + Q). \quad \text{(S1b)}$$

$\mathcal{S}(x)$, is a sigmoid function given by the equation:

$$\mathcal{S}(x) = \frac{1}{1 + e^{-a(x-\theta)}} - \frac{1}{1 + e^{a\theta}}, \quad (2)$$

With gain $a$ and threshold $\theta$. The form of the sigmoid in Equation S2 has been taken from the original formulation (Wilson and Cowan, 1972). Barring a missing negative sign before the gain '$a$' in the denominator of the equation provided in Supplementary C of (Buxton, 2021), a possible typing error, the two sigmoid functions are identical.

By virtue of these assumptions, the self-inhibitory activity and refractory dynamics were ignored, and the strength of all connections was assumed to be the same. Equations S1a and S1b were then rewritten as

$$\tau_0 \frac{dE}{dt} = -E + \mathcal{S}(wE - wI + P), \tag{S3a}$$

$$\tau_0 \frac{dI}{dt} = -I + \mathcal{S}(wE + Q). \tag{S3b}$$

wherein, $w = 3$, $\tau_0 = 10$ ms, $Q = 0.2$ and $P$ ranged from 0 to 2.5 (Buxton, 2021). Additionally, an initial rest condition was assumed: $E(0) = 0, I(0) = 0$. All subsequent steps were identical to the Buxton paper for the positive R
